# Older adults demonstrate interlimb transfer of reactive gait adaptations to repeated unpredictable gait perturbations

**DOI:** 10.1101/673574

**Authors:** Christopher McCrum, Kiros Karamanidis, Lotte Grevendonk, Wiebren Zijlstra, Kenneth Meijer

## Abstract

The ability to rapidly adjust gait to cope with unexpected mechanical perturbations declines with ageing. Previous studies however, have not ensured that pre-perturbation gait stability was equivalent, meaning that differences in unperturbed gait stability may have influenced the outcomes, which this study addresses. We also examine if interlimb transfer of gait adaptations are observed in healthy older adults, potentially driven by the increased motor error experienced due to their reduced ability to cope with the perturbations. 30 young and 28 older healthy adults experienced ten unpredictable treadmill belt accelerations (the first and last applied to the right leg, the others to the left) during walking at their stability-normalised walking speeds (young: 1.32±0.07m/s; older: 1.31±0.13m/s). Using kinematic data, we assessed the margins of stability during unperturbed walking and the first eight post-perturbation recovery steps. Older adults required three more steps to recover during the first perturbation to each leg than the young adults. Yet, after repeated perturbations of the left leg, older adults required only one more step to recover. Interestingly, for the untrained right leg, we found an improvement of three steps in the recovery of the older adults, indicating interlimb transfer of the improvements. Age differences in reactive gait stability remain even when participants’ walk with equivalent stability. Furthermore, we show that healthy older adults can transfer improvements in balance recovery made during repeated perturbations to one limb to their recovery following a perturbation to the untrained limb.

## Introduction

The ability to maintain or recover gait stability following unexpected mechanical perturbations such as trips, slips or ground surface changes is negatively affected in older age ^1-3^ which reflects older adults’ increased falls risk during walking ^4,5^. However, ageing does not greatly affect the ability to improve balance recovery responses to unexpected perturbations with repeated practice, nor the retention of these improvements over time ^6-9^. This has led to the development of perturbation-based balance training interventions, whereby different perturbations can be experienced and used as a training stimulus in a safe, controlled environment ^10^.

There is reasonable evidence in humans that increasing motor error during locomotion facilitates greater or faster adaptation ^11,12^. Motor error during a gait-like task in a stepping robot improves motor learning in young adults ^13,14^ and adaptation to split-belt and force-field perturbations during walking, as well as subsequent retention and savings of these adaptations, often occur to a greater extent following abrupt versus gradual exposure to the perturbations ^15-17^. Additionally, experiencing large, rather than small, perturbations in an initial task leads to better generalisation to other similar locomotor tasks in both split-belt walking ^18^ and slip-perturbed walking ^19^. These results indicate that older adults, with their reduced ability to maintain and recover balance following gait perturbations, may experience larger motor errors and experience a relatively larger stimulus for adaptation than younger adults completing the same gait perturbation task. In previous gait perturbation studies, transfer between similar perturbation tasks has been observed ^20-22^ but there is only limited evidence of interlimb transfer of reactive gait adaptations to perturbations ^23,24^. However, in both previous interlimb studies, only young healthy participants were included, meaning that the extent of motor errors experienced may have been much less than those that might be experienced by older adults under similar perturbation conditions.

Walking speed can influence the impact of, and the response to, different perturbations ^25-27^ and if the same speed is used for all participants, this may result in different degrees of task difficulty ^28,29^. In this study, we first aimed to determine if age-related differences in reactive gait stability and its adaptability in response to repeated mechanical gait perturbations are found when the participants’ walking speed is normalised to gait stability. To achieve this, we applied our recently published method of walking speed normalisation which reduces inter-participant differences in gait stability ^29^ assessed by the margins of stability (MoS) ^30^. With this method, an equivalent baseline level of gait stability across participants is achieved and we can infer that any differences in the response to the perturbations are not, in whole or part, artefacts of the walking speeds of the participants, but rather due to true differences in recovery responses. Based on previous work in trips leading to forward balance loss ^1,8^, we hypothesised that older adults would require more steps to recover stability than younger adults following the initial perturbation to each leg, despite the stability-normalised walking speed, but that both groups would be able to adapt their gait to improve stability over the repetitions to the left leg. The second aim was to determine if interlimb transfer of these adaptations could be observed in healthy older adults, despite the limited support in young adults in our previous study ^24^. Given our expectation that older adults would require more steps to recover stability than younger adults following the initial perturbation to each leg, and thereby experience greater motor error in their early responses, we hypothesised that evidence of interlimb transfer would be found in the older adults.

## Methods

### Participants

30 healthy young adults (12 males, 18 females; age: 24±2.5y; height: 173±8cm; weight: 71±13.9kg) and 28 healthy older adults (17 males, 11 females; age: 71±4y; height: 169±9.3cm; weight: 76±11.9kg) participated in this study. Participants were recruited via posters placed around the university and in local gyms and fitness centres. Data from 18 healthy young adults have been reported in our previous study ^24^ as part of a different analysis. While this was a convenience sample taken from a larger study powered for a different outcome, 28 to 30 participants provides sufficient power (0.72<β<0.96) to detect the moderate to large effect sizes of interest (Cohen’s d of 0.5-0.7) that we observed in our previous study ^24^. The participants had no self-reported history of walking difficulties, dizziness or balance problems, had no known neuromuscular condition or injury that could affect balance or walking, and could walk at a regular pace for 30 minutes without assistance and without stopping. Written informed consent was obtained and the study was conducted in accordance with the Declaration of Helsinki. The study protocol was approved by the Maastricht University Medical Centre medical ethics committee.

### Setup and Procedures

A dual-belt force plate-instrumented (1000Hz) treadmill with a virtual environment that provided optic flow during walking (Computer Assisted Rehabilitation Environment Extended, CAREN; Motekforce Link, Amsterdam, The Netherlands) and a 12-camera motion capture system (100Hz; Vicon Motion Systems, Oxford, UK) were used in this study. Three high definition video cameras recorded video footage of the trials. Five retroreflective markers were attached to anatomical landmarks (C7, left and right trochanter and left and right hallux) and the three-dimensional coordinates of these markers were tracked by the motion capture system. Participants were secured in a safety harness system throughout the measurements.

Participants first completed 60-second-long walking familiarisation trials at speeds of 0.4m/s up to 1.8m/s in 0.2m/s intervals and were given sufficient rest (approximately two minutes) before continuing with the recorded trials, comprised of two-to-three-minute-long trials at the same speeds. While participants rested, the stability-normalised walking speed was calculated by fitting a second order polynomial function to the mean anteroposterior margins of stability (MoS; see below) of the final 10 steps of each walking trial (0.4m/s to 1.8m/s) ^24,29^. The theoretical background and data on the effectiveness of this approach are described elsewhere ^29^. For each participant, the walking speed that would result in MoS of 0.05m was calculated from the function. The perturbation trial then began with three to four minutes of unperturbed walking at the stability-normalised walking speed, followed by 10 unannounced unilateral treadmill belt acceleration perturbations, each occurring every 30-90 seconds. Participants were told that they would complete a walking balance challenge and to try to continue walking as normally as possible. Participants were not aware of the specifics of the protocol (i.e. limbs to be perturbed, type, number, timing, magnitude of the perturbations). The first and tenth accelerations perturbed the right leg, while the second to ninth accelerations perturbed the left leg. The perturbation was a 3m/s^2^ acceleration of the treadmill belt to a maximum speed equal to 180% of the stability-normalised walking speed. The acceleration began when the hallux marker of the to-be-perturbed limb passed the hallux marker of the opposite foot in the sagittal plane. The belt decelerated at toe-off of the perturbed limb.

### Data Processing and Margin of Stability Calculation

Data processing was conducted in MATLAB (2016a, The MathWorks, Inc., Natick). The three-dimensional coordinates of the markers were filtered using a low pass second order Butterworth filter (zero-phase) with a 12Hz cut-off frequency. Foot touchdown and toe-off were determined as previously described ^24,29^. The anteroposterior MoS (MoS_AP_) at foot touchdown were calculated as the anteroposterior distance between the anterior boundary of the base of support (BoS) and the extrapolated centre of mass (X_CoM_) ^30^, adapted for our reduced kinematic model ^1^. The mediolateral MoS (MoS_ML_) were also calculated in a similar manner (mediolateral components instead of anteroposterior), with the exceptions that the treadmill belt velocity was not included in the estimation of CoM velocity and that the MoS_ML_ was not determined at foot touchdown, but rather the minimum MoS_ML_ during the stance phase was determined ^31^. The MoS was calculated for the following steps: baseline for each perturbation was the mean MoS of the eleventh to second last step before each perturbation (Base); the final step before each perturbation (Pre); and the first eight recovery steps following each perturbation (Post1-8).

### Statistics

Two-way repeated measures ANOVAs with group (young, older) and step (Base, Pre, Post1-Post8) as factors were conducted individually for the first, second and ninth perturbations (the first perturbation of the right leg and the first and final perturbations of the left “trained” leg; Pert1_R_, Pert2_L_ and Pert9_L_ respectively). To evaluate any changes in the MoS_AP_ during unperturbed walking, a repeated measure mixed model with perturbation number and age group as factors was conducted. To further investigate which components might be responsible for any observed differences, the same ANOVAs were conducted for the BoS and X_CoM_. Finally, the same ANOVAs were conducted for the MoS_ML_, as we suspected that lateral instability may also be increased in the older adults during anteroposterior perturbations ^32^. To determine if interlimb transfer of the reactive adaptations occurred in the older adults, two-way repeated measures ANOVAs with perturbation number (Pert1_R_ and Pert10_R_) and step (Base, Pre, Post1-Post8) as factors were conducted for MoS_AP_, BoS and X_CoM_. For all ANOVAs, post hoc Bonferroni tests for multiple comparisons were applied. Sphericity of the data was checked and when required, outcomes were adjusted using the Geisser-Greenhouse epsilon. Significance was set at α=0.05. Analyses were performed using GraphPad Prism version 8.02 for Windows (GraphPad Software Inc., La Jolla, California, USA).

## Results

Similar to our previous work in young adults ^24,29^, means and SDs of the eleventh to second last step before the first perturbation revealed that most participants (25 of 30 young adults and 23 of 28 older adults) were within one SD of the desired 0.05m MoS_AP_ (Fig. 1). The stability-normalised walking speeds (mean±SD, range) were 1.32±0.07m/s, 1.16-1.51m/s for the young adults and 1.31±0.13m/s, 1.01-1.50m/s for the older adults.

**Fig 1:**
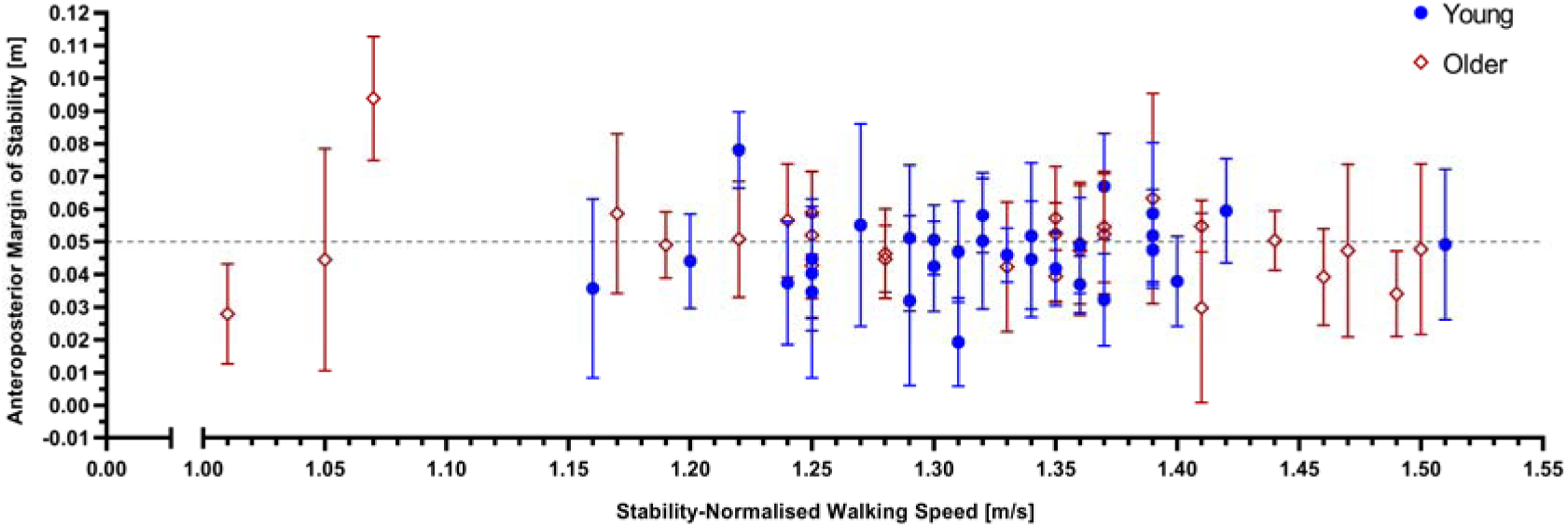
Anteroposterior margins of stability (means and SDs) of the eleventh to second last step before the first perturbation across the individual stability-normalised walking speeds for young (blue circles) and older (red diamonds) healthy adults.

All participants completed the task without harness assistance. However, one older adult stopped walking after recovering from the first perturbation, leading to the treadmill stopping. This participant was therefore excluded from the analyses involving Pert1_R_. Two way repeated measures ANOVAs for Pert1_R_, Pert2_L_ and Pert9_L_ revealed a significant age group effect on MoS_AP_ for Pert1_R_ only (Pert1_R_: F_(1, 55)_=14.11, P=0.0004; Pert2_L_: F_(1, 56)_=2.968, P=0.0904; Pert9_L_: F_(1, 56)_=0.2948, P=0.5893). Significant Step by Age Group interactions were found for Pert1_R_ and Pert2_L_ (Pert1_R_: F_(9, 495)_=15.55, P<0.0001; Pert2_L_: F_(9, 504)_=8.310, P<0.0001; Pert9_L_: F_(9, 504)_=1.576, P=0.1192). Bonferroni tests for multiple comparisons (indicated in Fig. 2) revealed that, on average, older adults had MoS_AP_ significantly different to Base for at least three steps more than the young adults during Pert1_R_ and Pert2_L_, but during Pert9_L_ older adults had MoS_AP_ significantly different to Base for only one step more than the young adults (five vs. four steps). The young and older adults improved their recovery performance following repeated perturbations (Pert2_L_ to Pert9_L_) on average by two and three recovery steps, respectively. Complete Bonferroni results can be found in the supplement (eTables 1 and 2). Regarding the unperturbed walking MoS_AP_, we did find a significant perturbation number effect (F_(3, 166)_=11.44, P<0.0001) and Bonferroni post hoc tests revealed significant differences between Pert1_R_ and Pert9_L_, Pert2_L_ and Pert9_L_ and between Pert2_L_ and Pert10_R_ in the younger adults and between Pert2_L_ and Pert9_L_ in the older adults, but these differences ranged from 0.2cm to 0.8cm and were therefore not considered to have a meaningful effect on the main results.

**Fig. 2:**
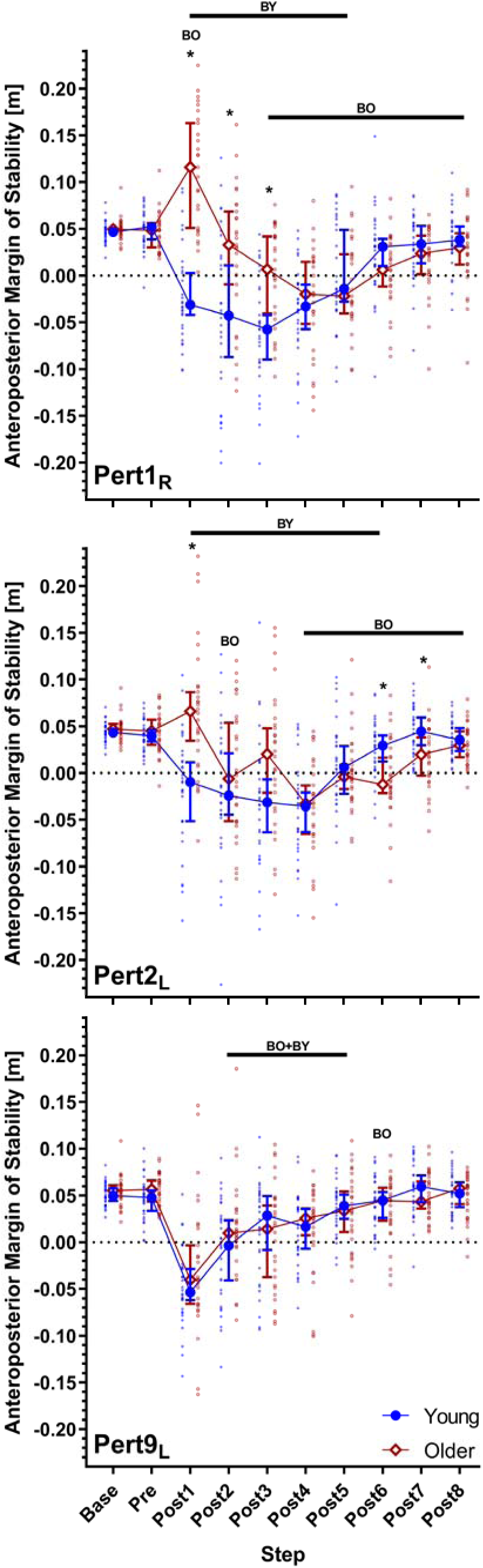
Median and 95% confidence intervals (with individual data points) of the anteroposterior margins of stability during the first, second and ninth perturbations (Pert1_R_, Pert2_L_, and Pert9_L_, respectively) including unperturbed walking prior to each perturbation (Base), the final step prior to each perturbation (Pre) and the first eight recovery steps following the perturbations (Post1-8) for young and older adults. BO and BY: Significant difference to Base for older and young adults, respectively (P<0.05). *: Significant difference between young and older adults (P<0.05).

Two way repeated measures ANOVAs for Pert1_R_, Pert2_L_ and Pert9_L_ revealed significant age group effects on BoS for Pert1_R_, Pert2_L_ and Pert9_L_ (Pert1_R_: F_(1, 55)_=7.862, P=0.007; Pert2_L_: F_(1, 56)_=11.75, P=0.0011; Pert9_L_: F_(1, 56)_=9.078, P=0.0039; Fig. 3). Significant Step by Age Group interactions were found for Pert1_R_, Pert2_L_ and Pert9_L_ (Pert1_R_: F_(9, 495)_=3.160, P=0.001; Pert2_L_: F_(9, 504)_=7.281, P<0.0001; Pert9_L_: F_(9, 504)_=1.987, P=0.0389; Fig. 3). Bonferroni tests for multiple comparisons (indicated in Fig. 3) revealed that, on average, older adults had returned to BoS values not significantly different to Base by Post4 during each of the analysed perturbations. Older adults had a significantly smaller BoS than young adults during Post2 to Post4 for Pert2_L_, and Post3 and Post4 for Pert9_L_. For X_CoM_, the ANOVAs revealed significant age group effects (Pert1_R_: F_(1, 55)_=16.26, P=0.0002; Pert2_L_: F_(1,_ 56)=15.64, P=0.0002; Pert9_L_: F_(1, 56)_=9.140, P=0.0038; Fig. 3) and Step by Age Group interactions (Pert1_R_: F_(9, 495)_=10.45, P<0.0001; Pert2_L_: F_(9, 504)_=11.84, P<0.0001; Pert9_L_: F_(9,_ 504)=2.440, P=0.0101; Fig. 3) for Pert1_R_, Pert2_L_ and Pert9_L_. Bonferroni tests for multiple comparisons revealed that X_CoM_ significantly differed between older and young adults from Post1 to Post4 for Pert1_R_ and Pert2_L_ (Fig. 3). Complete Bonferroni results for the BoS and X_CoM_ can be found in the supplement (eTables 3 to 6). Results regarding the MoS_ML_ can be found in the supplement (eResults, eFigure 1, eTables 7 and 8)

**Fig. 3:**
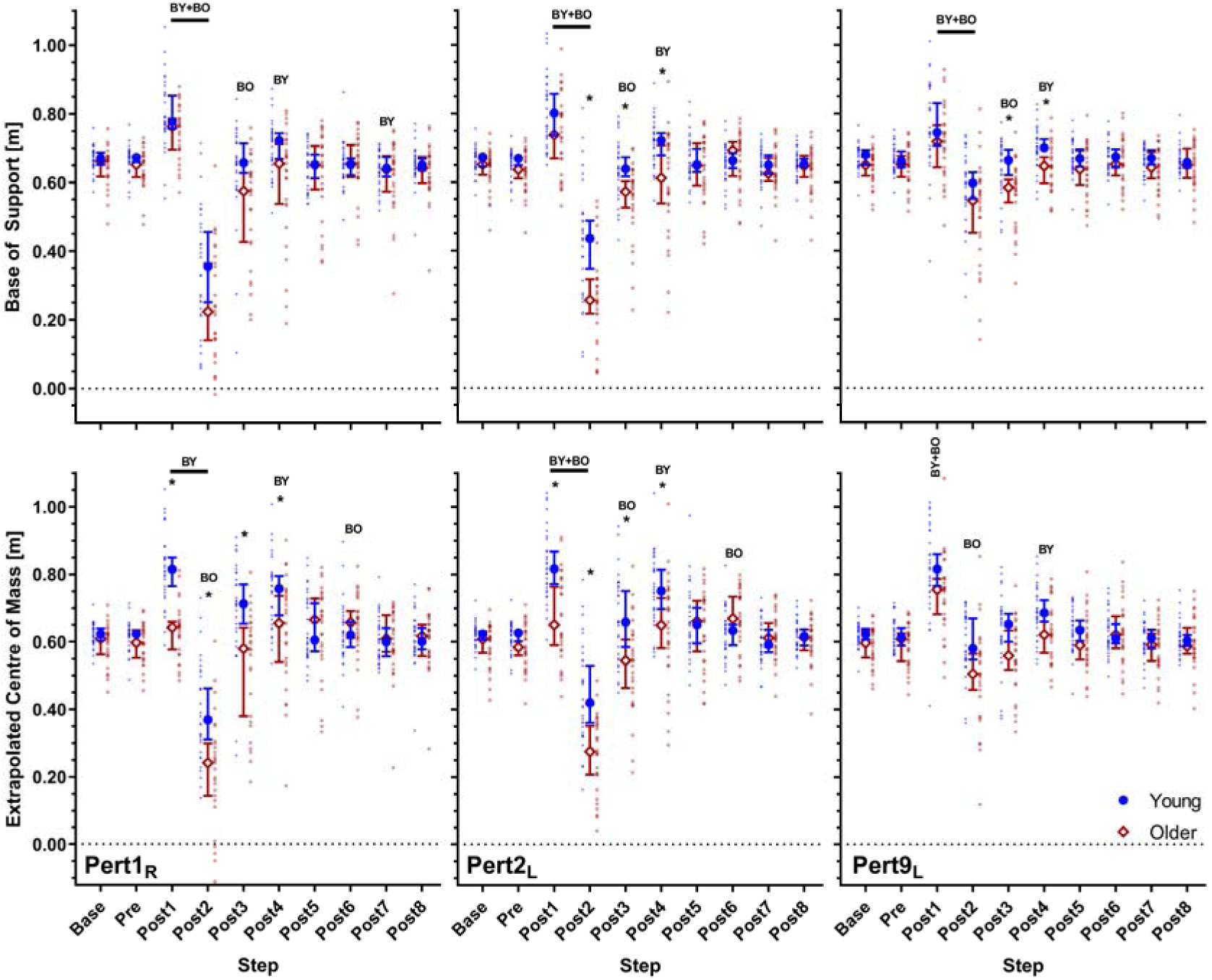
Median and 95% confidence intervals (with individual data points) of the anteroposterior base of support and extrapolated centre of mass during the first, second and ninth perturbations (Pert1_R_, Pert2_L_, and Pert9_L_, respectively) including unperturbed walking prior to each perturbation (Base), the final step prior to each perturbation (Pre) and the first eight recovery steps following the perturbations (Post1-8) for young and older adults. BO and BY: Significant difference to Base for older and young adults, respectively (P<0.05). *: Significant difference between young and older adults (P<0.05).

Regarding the investigation of interlimb transfer in the older adults (Pert1_R_ and Pert10_R_), no significant perturbation number effects were found for MoS_AP_ or MoS_ML_ (MoS_AP_: F_(1, 26)_=2.634, P=0.1167; MoS_ML_: F_(1, 26)_= 0.03025, P=0.8633; Fig. 5). However, significant perturbation number effects were found for BoS and X_CoM_ (BoS: F_(1, 26)_=9.104, P=0.0056; X_CoM_: F_(1, 26)_=18.32, P=0.0002; Fig. 5), along with significant perturbation number by step interactions for MoS_AP_, BoS and X_CoM_, but not MoS_ML_ (MoS_AP_: F_(4.150,_ 107.9)=16.42, 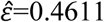, P<0.0001; BoS: F_(3.029, 78.74)_=5.480, 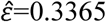, P=0.0017; X_CoM_: F_(3.920,_ 101.9)=12.30, 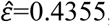, P<0.0001; MoS_ML_: F_(4.056, 105.5)_=0.6885, 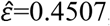, P=0.6035; Fig. 5).

Bonferroni tests for multiple comparisons are indicated in Fig. 4 and revealed that during Pert1_R_, the older adults did not return to MoS_AP_ values not significantly different to Base during the eight analysed recovery steps, whereas during Pert10_R_, they reached this point by Post6. During Pert1_R_, Post1 and Post2 had significantly greater MoS_AP_ than during Pert10_R_, but significantly lower MoS_AP_ during Post8. BoS was significantly smaller during Pert1_R_ than Pert10_R_ at Post2. This coincided with significant differences between Pert1_R_ and Pert10_R_ in X_CoM_ at Post1, Post2 and Post3, with more anterior X_CoM_ during Pert10_R_. No significant differences in MoS_ML_ were observed between Pert1_R_ and Pert10_R_. Complete Bonferroni results for the examination of interlimb transfer can be found in the supplement (eTables 9 to 16).

**Fig. 4:**
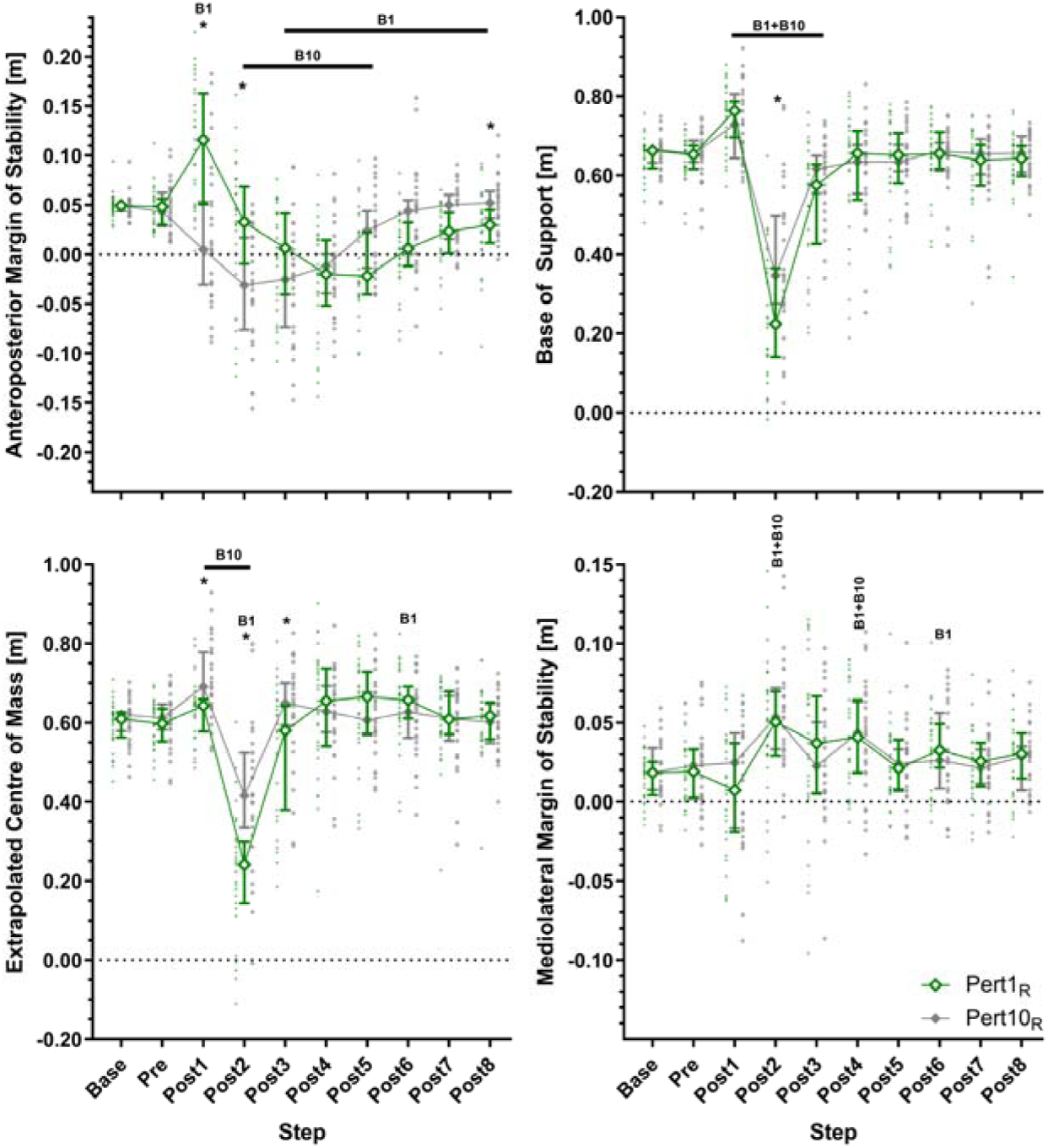
Median and 95% confidence intervals (with individual data points) of the anteroposterior margins of stability, base of support, extrapolated centre of mass and mediolateral margins of stability during the first and tenth perturbations (the first and final perturbations to the right leg; Pert1_R_ and Pert10_R_, respectively) including unperturbed walking prior to each perturbation (Base), the final step prior to each perturbation (Pre) and the first eight recovery steps following the perturbations (Post1-8) for older adults. B1 and B10: Significant difference to Base for Pert1_R_ and Pert10_R_, respectively (P<0.05). *: Significant difference between Pert1_R_ and Pert10_R_ (P<0.05).

## Discussion

The current study aimed to determine if age-related differences in reactive gait stability and its adaptability in response to repeated mechanical gait perturbations are found when the participants’ walking speed is normalised to gait stability, and if evidence of interlimb transfer of these adaptations can be observed in healthy older adults. We hypothesised that older adults require more steps to recover stability than younger adults following the initial perturbation to each leg, despite the stability-normalised walking speed, but that both groups would be able to adapt their gait to improve stability over the repetitions to the left leg. These hypotheses were confirmed, as the older adults required approximately three more steps to recover stability during the first perturbations to each leg than the young adults, but after repeated perturbations of the left leg, required approximately three fewer steps to recover than during the first perturbations and no longer showed significant differences to the young adults at any recovery step. These findings confirm previous studies in older adults using slip, trip and surface compliance perturbations ^2,6-9,33^ and extends these to treadmill belt acceleration perturbations during which the walking speed is normalised to stability and the perturbation is normalised to the walking speed, ensuring equivalent baseline gait stability and task difficulty. We also hypothesised that evidence of interlimb transfer would be found in the older adults due to them experiencing greater motor error in their early responses. This hypothesis was confirmed, as we found a three-step improvement in the steps to reach MoS_AP_ values not significantly different to Base, as well as a more anterior X_CoM_ position during Post1 to Post3 in Pert10_R_ compared to Pert1_R_.

Ageing has repeatedly been shown to be associated with poorer performance in recovering stability following unexpected gait perturbations ^1-3^. However, as previously described, potential differences in gait as a result of the walking speed choices in previous studies may have affected these findings ^28,29^. The current study confirms and consolidates previously reported age-related differences in reactive gait stability, as age differences were observed despite the use of individual stability-normalised walking speeds. We found that these age differences in MoS_AP_ were the result of significantly smaller X_CoM_ during the first four recovery steps following the first two perturbations and to a lesser extent, smaller BoS during the second to fourth recovery steps following the first and second perturbations. These results indicate that the older adults responded to the treadmill belt acceleration perturbation with a more posterior X_CoM_ and smaller BoS than young adults, delaying their stability recovery. This differs to what we have previously observed using a cable-trip setup, where the differences have been observed in the BoS ^8,34,35^, reflecting the differences in perturbation type. By the final perturbation of the trained leg, more posterior X_CoM_ (not significant) and smaller BoS values were still visible in the older adults compared to the young adults, but these no longer led to significantly different MoS_AP_ values. Multiple studies have demonstrated the ability of healthy adults to reactively adapt gait in response to repeated perturbations ^6-9^ and the current study confirms these findings in a treadmill belt acceleration paradigm with stability-normalised walking speeds and walking speed normalised perturbations. Therefore, we can conclude that potential differences in the initial gait stability or perturbation characteristics likely do not play a large role in whether older adults adapt their response to repeated perturbations.

We previously found little support in young adults for interlimb transfer of reactive gait adaptations following the same protocol as the current study ^24^. However, we expected that older adults would require more steps to recover stability than younger adults following the initial perturbation to each leg, and thereby they would experience greater motor error in their early responses that may stimulate interlimb transfer. Our results confirmed this expectation, as we found a three-step improvement in the steps to reach MoS_AP_ values not significantly different to Base from Pert1_R_ to Pert10_R_. In addition, perturbation number effects on BoS and X_CoM_ were found. The older adults appeared to respond to Pert1_R_ with a more posterior X_CoM_ at Post1 than in Pert10_R_, and with a smaller BoS and posterior X_CoM_ at Post2. This alteration in recovery strategy during Post1-3 resulted in the three-step reduction in reaching MoS_AP_ values not significantly different to Base. Therefore, it appears that both the overall recovery performance and the altered movement strategy were transferred to the untrained leg.

An interesting finding of the current study was that the older adults during Pert1_R_ demonstrated an increase, rather than a decrease in stability at Post1, whereas the young adults during all perturbations and the older adults during Pert9_L_ and Pert10_R_ (Figs. 2 and 4) demonstrated a decrease in stability. This increase was caused by a more posterior X_CoM_ during Post1 in Pert1_R_, but not a difference in BoS, implying that trunk motion was at least partly responsible. Future work could further investigate this using a kinematic model more suited to assessing trunk motion in detail. We speculate that this may be one potential reason for the observed interlimb transfer of balance recovery performance. While the lower limbs may play very specific roles in perturbation recovery during the first recovery step (i.e. push-off versus swing and placing of the foot), the role of the trunk may be more generalisable across perturbations to different limbs (i.e. counter-rotation to forward balance loss). This may also explain why in our previous article, no clear interlimb transfer occurred ^24^, because young adults do not show as posterior an X_CoM_ position (Fig. 3). Regarding our analyses of MoS_ML_, the results did not reveal any substantial differences with age, and these are discussed in the supplement (eDiscussion).

A limitation of the current work is that it is unclear if these findings would generalise to populations with reduced locomotor function and it is these groups that potentially could benefit most from perturbation-based balance training programmes ^10^. Therefore, interlimb and intertask transfer of adaptations in reactive balance control and the generalisability of these improvements to daily life should be further explored. It could be argued that leg dominance may have affected the results, but due to the bipedal nature of the task, we think this is unlikely. Only one study has specifically investigated the effect of limb dominance on recovery from sudden balance loss and found no differences in performance between stepping with the dominant and non-dominant limbs in young and older adults ^36^. Another limitation worth considering is that despite the evidence provided here that interlimb transfer can occur during a single short perturbation session, this does not necessarily imply that this will be retained over time, as perturbation dose appears to be related to the degree of retention possible ^37^.

In conclusion, the current results show that healthy older adults have a decreased ability to cope with unpredictable gait perturbations compared with younger adults, even when their walking speeds are normalised to an equivalent stability value. However, as previous studies have also shown, older healthy adults are capable of reactively adapting their gait to improve their stability following repeated gait perturbations and can then perform similarly to young adults. The current study provides evidence that older adults can transfer improvements in the number of steps required for balance recovery following repeated perturbations to one limb to their recovery following a perturbation to the untrained limb, which in this study was mostly due to an alteration in the X_CoM_ position, rather than in the BoS.

## Acknowledgements

The authors thank Paul Willems, Wouter Bijnens and Marissa Gerards for their support during this project.

