## Supplement for "Older adults demonstrate interlimb transfer of reactive gait adaptations to repeated unpredictable gait perturbations"

### eResults

Regarding the $MoS_{ML}$, the two way repeated measures ANOVAs for $Pert1_R$, $Pert2_L$ and $Pert9_L$ revealed a significant age effect for $Pert1_R$ and $Pert9_L$, but not for $Pert2_L$ ($Pert1_R$: $F_{(1, 55)}$=4.973, P=0.0298; $Pert2_L$: $F_{(1, 56)}$=2.031, P=0.1597; $Pert9_L$: $F_{(1, 56)}$=4.110, P=0.0474; Fig. 4). Significant Age Group by Step interactions were found for $Pert1_R$, $Pert2_L$ and $Pert9_L$ ($Pert1_R$: $F_{(9, 495)}$=1.965, P=0.0415; $Pert2_L$: $F_{(9, 504)}$=2.254, P=0.0177; $Pert9_L$: $F_{(9, 504)}$=3.610, P=0.0002; Fig. 4). Bonferroni tests for multiple comparisons are indicated in Fig. 4 and revealed two significant difference between the age groups at Post4 during $Pert1_R$, and significant differences at Post5 and Post6 during $Pert9_L$ but no major differences were identified pre and post repetition of the left limb perturbations, with both age groups showing significantly increased $MoS_{ML}$ at Post2 and Post3 during $Pert2_L$ and $Pert9_L$. Complete Bonferroni results can be found in Supplementary Tables 7 and 8.

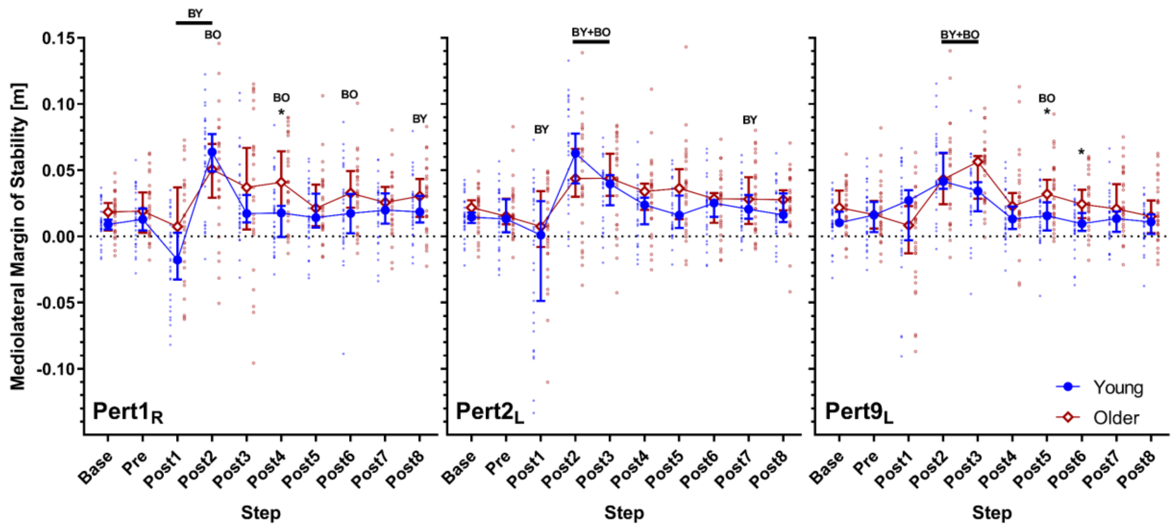


**eFigure 1**: Median and 95% confidence intervals (with individual data points) of the mediolateral margin of stability during the first, second and ninth perturbations ($Pert1_R$, $Pert2_L$, and $Pert9_L$, respectively) including unperturbed walking prior to each perturbation (Base), the final step prior to each perturbation (Pre) and the first eight recovery steps following the perturbations (Post1-8) for young and older adults. BO and BY: Significant difference to Base for older and young adults, respectively (P<0.05). *: Significant difference between young and older adults (P<0.05).

**eTable 1: Bonferroni's multiple comparison test results comparing recovery steps to baseline in $Pert1_R$, $Pert2_L$ and $Pert9_L$ for $MoS_{AP}$ in the young and older adults**

| Perturbation | Test details | Mean 1 | Mean 2 | Mean Diff. | SE of diff. | N1 | N2 | t | DF | 95.00% CI of diff. | Adjusted P Value |
|---|---|---|---|---|---|---|---|---|---|---|---|
| **$Pert1_R$** | Young | | | | | | | | | | |
| | Base vs. Pre | 0.04649 | 0.04944 | -0.002954 | 0.003325 | 30 | 30 | 0.8884 | 29 | -0.01292 to 0.007008 | >0.9999 |
| | Base vs. Post1 | 0.04649 | -0.01941 | 0.0659 | 0.009999 | 30 | 30 | 6.59 | 29 | 0.03594 to 0.09585 | <0.0001 |
| | Base vs. Post2 | 0.04649 | -0.04436 | 0.09085 | 0.01635 | 30 | 30 | 5.556 | 29 | 0.04186 to 0.1398 | <0.0001 |
| | Base vs. Post3 | 0.04649 | -0.06068 | 0.1072 | 0.01102 | 30 | 30 | 9.721 | 29 | 0.07414 to 0.1402 | <0.0001 |
| | Base vs. Post4 | 0.04649 | -0.03958 | 0.08607 | 0.009342 | 30 | 30 | 9.213 | 29 | 0.05808 to 0.1141 | <0.0001 |
| | Base vs. Post5 | 0.04649 | 0.00193 | 0.04456 | 0.0101 | 30 | 30 | 4.411 | 29 | 0.01429 to 0.07482 | 0.0012 |
| | Base vs. Post6 | 0.04649 | 0.02455 | 0.02194 | 0.008469 | 30 | 30 | 2.591 | 29 | -0.003432 to 0.04732 | 0.1335 |
| | Base vs. Post7 | 0.04649 | 0.03039 | 0.0161 | 0.006845 | 30 | 30 | 2.351 | 29 | -0.004411 to 0.03660 | 0.2313 |
| | Base vs. Post8 | 0.04649 | 0.03976 | 0.006733 | 0.005854 | 30 | 30 | 1.15 | 29 | -0.01080 to 0.02427 | >0.9999 |
| | Older | | | | | | | | | | |
| | Base vs. Pre | 0.0496 | 0.04576 | 0.003836 | 0.003399 | 27 | 27 | 1.128 | 26 | -0.006443 to 0.01411 | >0.9999 |
| | Base vs. Post1 | 0.0496 | 0.1117 | -0.06213 | 0.01244 | 27 | 27 | 4.996 | 26 | -0.09973 to -0.02452 | 0.0003 |
| | Base vs. Post2 | 0.0496 | 0.02267 | 0.02693 | 0.01389 | 27 | 27 | 1.939 | 26 | -0.01506 to 0.06892 | 0.5709 |
| | Base vs. Post3 | 0.0496 | -0.0007875 | 0.05038 | 0.01063 | 27 | 27 | 4.739 | 26 | 0.01824 to 0.08253 | 0.0006 |
| | Base vs. Post4 | 0.0496 | -0.02787 | 0.07746 | 0.01084 | 27 | 27 | 7.143 | 26 | 0.04467 to 0.1103 | <0.0001 |
| | Base vs. Post5 | 0.0496 | -0.009746 | 0.05934 | 0.01015 | 27 | 27 | 5.846 | 26 | 0.02865 to 0.09004 | <0.0001 |
| | Base vs. Post6 | 0.0496 | 0.004204 | 0.04539 | 0.007151 | 27 | 27 | 6.348 | 26 | 0.02377 to 0.06702 | <0.0001 |
| | Base vs. Post7 | 0.0496 | 0.01565 | 0.03395 | 0.007473 | 27 | 27 | 4.542 | 26 | 0.01135 to 0.05654 | 0.001 |
| | Base vs. Post8 | 0.0496 | 0.02613 | 0.02347 | 0.0077 | 27 | 27 | 3.048 | 26 | 0.0001856 to 0.04675 | 0.0471 |
| **$Pert2_L$** | Young | | | | | | | | | | |
| | Base vs. Pre | 0.04342 | 0.03979 | 0.003637 | 0.002281 | 30 | 30 | 1.594 | 29 | -0.003198 to 0.01047 | >0.9999 |
| | Base vs. Post1 | 0.04342 | -0.01938 | 0.06281 | 0.01185 | 30 | 30 | 5.299 | 29 | 0.02730 to 0.09831 | <0.0001 |
| | Base vs. Post2 | 0.04342 | -0.02011 | 0.06353 | 0.01373 | 30 | 30 | 4.626 | 29 | 0.02239 to 0.1047 | 0.0006 |
| | Base vs. Post3 | 0.04342 | -0.03096 | 0.07438 | 0.01267 | 30 | 30 | 5.87 | 29 | 0.03642 to 0.1123 | <0.0001 |
| | Base vs. Post4 | 0.04342 | -0.03964 | 0.08307 | 0.009214 | 30 | 30 | 9.016 | 29 | 0.05546 to 0.1107 | <0.0001 |
| | Base vs. Post5 | 0.04342 | 0.00629 | 0.03713 | 0.009356 | 30 | 30 | 3.969 | 29 | 0.009103 to 0.06516 | 0.0039 |
| | Base vs. Post6 | 0.04342 | 0.0266 | 0.01682 | 0.005298 | 30 | 30 | 3.175 | 29 | 0.0009481 to 0.03269 | 0.0318 |
| | Base vs. Post7 | 0.04342 | 0.04451 | -0.001082 | 0.004687 | 30 | 30 | 0.2309 | 29 | -0.01512 to 0.01296 | >0.9999 |
| | Base vs. Post8 | 0.04342 | 0.0363 | 0.007127 | 0.003821 | 30 | 30 | 1.865 | 29 | -0.004321 to 0.01857 | 0.6509 |
| | Older | | | | | | | | | | |
| | Base vs. Pre | 0.0478 | 0.04399 | 0.003813 | 0.004662 | 28 | 28 | 0.8179 | 27 | -0.01024 to 0.01786 | >0.9999 |
| | Base vs. Post1 | 0.0478 | 0.06765 | -0.01985 | 0.01443 | 28 | 28 | 1.375 | 27 | -0.06334 to 0.02365 | >0.9999 |
| | Base vs. Post2 | 0.0478 | 0.002104 | 0.0457 | 0.01349 | 28 | 28 | 3.388 | 27 | 0.005044 to 0.08635 | 0.0196 |
| | Base vs. Post3 | 0.0478 | 0.01392 | 0.03388 | 0.01371 | 28 | 28 | 2.471 | 27 | -0.007437 to 0.07520 | 0.1806 |
| | Base vs. Post4 | 0.0478 | -0.03748 | 0.08529 | 0.01072 | 28 | 28 | 7.954 | 27 | 0.05297 to 0.1176 | <0.0001 |

| | | | | | | | | | | | |
|---|---|---|---|---|---|---|---|---|---|---|---|
| | Base vs. Post5 | 0.0478 | 0.001129 | 0.04667 | 0.008095 | 28 | 28 | 5.765 | 27 | 0.02228 to 0.07107 | <0.0001 |
| | Base vs. Post6 | 0.0478 | -0.005296 | 0.0531 | 0.007842 | 28 | 28 | 6.771 | 27 | 0.02946 to 0.07673 | <0.0001 |
| | Base vs. Post7 | 0.0478 | 0.01784 | 0.02997 | 0.006037 | 28 | 28 | 4.964 | 27 | 0.01177 to 0.04816 | 0.0003 |
| | Base vs. Post8 | 0.0478 | 0.0303 | 0.0175 | 0.004551 | 28 | 28 | 3.845 | 27 | 0.003782 to 0.03121 | 0.006 |
| **Pert9$_L$** | Young | | | | | | | | | | |
| | Base vs. Pre | 0.05127 | 0.04608 | 0.005184 | 0.003708 | 30 | 30 | 1.398 | 29 | -0.005926 to 0.01629 | >0.9999 |
| | Base vs. Post1 | 0.05127 | -0.04829 | 0.09956 | 0.008349 | 30 | 30 | 11.92 | 29 | 0.07454 to 0.1246 | <0.0001 |
| | Base vs. Post2 | 0.05127 | -0.01154 | 0.06281 | 0.01071 | 30 | 30 | 5.864 | 29 | 0.03072 to 0.09490 | <0.0001 |
| | Base vs. Post3 | 0.05127 | 0.01292 | 0.03835 | 0.01047 | 30 | 30 | 3.663 | 29 | 0.006980 to 0.06972 | 0.0089 |
| | Base vs. Post4 | 0.05127 | 0.01281 | 0.03846 | 0.007465 | 30 | 30 | 5.151 | 29 | 0.01609 to 0.06082 | 0.0001 |
| | Base vs. Post5 | 0.05127 | 0.03754 | 0.01373 | 0.005653 | 30 | 30 | 2.429 | 29 | -0.003205 to 0.03067 | 0.1941 |
| | Base vs. Post6 | 0.05127 | 0.04199 | 0.009278 | 0.00488 | 30 | 30 | 1.901 | 29 | -0.005343 to 0.02390 | 0.6054 |
| | Base vs. Post7 | 0.05127 | 0.05671 | -0.005442 | 0.005333 | 30 | 30 | 1.02 | 29 | -0.02142 to 0.01053 | >0.9999 |
| | Base vs. Post8 | 0.05127 | 0.05212 | -0.0008489 | 0.003863 | 30 | 30 | 0.2198 | 29 | -0.01242 to 0.01072 | >0.9999 |
| | Older | | | | | | | | | | |
| | Base vs. Pre | 0.05456 | 0.05559 | -0.001034 | 0.0039 | 28 | 28 | 0.2651 | 27 | -0.01279 to 0.01072 | >0.9999 |
| | Base vs. Post1 | 0.05456 | -0.02543 | 0.07999 | 0.01349 | 28 | 28 | 5.93 | 27 | 0.03934 to 0.1206 | <0.0001 |
| | Base vs. Post2 | 0.05456 | 0.01246 | 0.0421 | 0.01043 | 28 | 28 | 4.038 | 27 | 0.01068 to 0.07352 | 0.0036 |
| | Base vs. Post3 | 0.05456 | 0.004749 | 0.04981 | 0.01065 | 28 | 28 | 4.679 | 27 | 0.01773 to 0.08189 | 0.0006 |
| | Base vs. Post4 | 0.05456 | 0.01211 | 0.04245 | 0.009851 | 28 | 28 | 4.309 | 27 | 0.01276 to 0.07214 | 0.0018 |
| | Base vs. Post5 | 0.05456 | 0.02975 | 0.02481 | 0.008151 | 28 | 28 | 3.044 | 27 | 0.0002450 to 0.04937 | 0.0464 |
| | Base vs. Post6 | 0.05456 | 0.03516 | 0.0194 | 0.006734 | 28 | 28 | 2.881 | 27 | -0.0008939 to 0.03969 | 0.0691 |
| | Base vs. Post7 | 0.05456 | 0.04977 | 0.004786 | 0.003699 | 28 | 28 | 1.294 | 27 | -0.006363 to 0.01594 | >0.9999 |
| | Base vs. Post8 | 0.05456 | 0.04842 | 0.006134 | 0.00423 | 28 | 28 | 1.45 | 27 | -0.006613 to 0.01888 | >0.9999 |

**eTable 2: Bonferroni's multiple comparison test results comparing the young and older adults in Pert1$_R$, Pert2$_L$ and Pert9$_L$ for MoS$_{AP}$**

| Perturbation | Test details | Mean 1 | Mean 2 | Mean Diff. | SE of diff. | N1 | N2 | t | DF | 95.00% CI of diff. | Adjusted P Value |
|---|---|---|---|---|---|---|---|---|---|---|---|
| **Pert1$_R$** | Young - Older | | | | | | | | | | |
| | Base | 0.04649 | 0.0496 | -0.003109 | 0.00317 | 30 | 27 | 0.9805 | 53.23 | -0.01239 to 0.006177 | >0.9999 |
| | Pre | 0.04944 | 0.04576 | 0.003681 | 0.005625 | 30 | 27 | 0.6545 | 54.19 | -0.01278 to 0.02014 | >0.9999 |
| | Post1 | -0.01941 | 0.1117 | -0.1311 | 0.01545 | 30 | 27 | 8.488 | 51.21 | -0.1765 to -0.08581 | <0.0001 |
| | Post2 | -0.04436 | 0.02267 | -0.06703 | 0.02173 | 30 | 27 | 3.085 | 54.55 | -0.1306 to -0.003460 | 0.0319 |
| | Post3 | -0.06068 | -0.0007875 | -0.0599 | 0.01492 | 30 | 27 | 4.014 | 54.99 | -0.1035 to -0.01626 | 0.0018 |
| | Post4 | -0.03958 | -0.02787 | -0.01171 | 0.01481 | 30 | 27 | 0.7908 | 52.49 | -0.05511 to 0.03169 | >0.9999 |
| | Post5 | 0.00193 | -0.009746 | 0.01168 | 0.01426 | 30 | 27 | 0.8191 | 54.99 | -0.03002 to 0.05337 | >0.9999 |
| | Post6 | 0.02455 | 0.004204 | 0.02034 | 0.01151 | 30 | 27 | 1.768 | 54.9 | -0.01331 to 0.05400 | 0.8262 |
| | Post7 | 0.03039 | 0.01565 | 0.01474 | 0.009696 | 30 | 27 | 1.52 | 54.19 | -0.01363 to 0.04312 | >0.9999 |
| | Post8 | 0.03976 | 0.02613 | 0.01363 | 0.009728 | 30 | 27 | 1.401 | 48.45 | -0.01499 to 0.04224 | >0.9999 |
| **Pert2$_L$** | Young - Older | | | | | | | | | | |
| | Base | 0.04342 | 0.0478 | -0.004378 | 0.003218 | 30 | 28 | 1.36 | 50.76 | -0.01382 to 0.005067 | >0.9999 |
| | Pre | 0.03979 | 0.04399 | -0.004202 | 0.005268 | 30 | 28 | 0.7978 | 49.01 | -0.01969 to 0.01128 | >0.9999 |
| | Post1 | -0.01938 | 0.06765 | -0.08703 | 0.01795 | 30 | 28 | 4.848 | 53.7 | -0.1396 to -0.03447 | 0.0001 |
| | Post2 | -0.02011 | 0.002104 | -0.02221 | 0.01918 | 30 | 28 | 1.158 | 55.98 | -0.07827 to 0.03384 | >0.9999 |
| | Post3 | -0.03096 | 0.01392 | -0.04488 | 0.01904 | 30 | 28 | 2.358 | 55.04 | -0.1006 to 0.01080 | 0.2198 |
| | Post4 | -0.03964 | -0.03748 | -0.002159 | 0.01369 | 30 | 28 | 0.1577 | 55.61 | -0.04219 to 0.03787 | >0.9999 |
| | Post5 | 0.00629 | 0.001129 | 0.005161 | 0.01246 | 30 | 28 | 0.4143 | 54.94 | -0.03127 to 0.04160 | >0.9999 |
| | Post6 | 0.0266 | -0.005296 | 0.0319 | 0.009763 | 30 | 28 | 3.267 | 48.6 | 0.003185 to 0.06061 | 0.02 |
| | Post7 | 0.04451 | 0.01784 | 0.02667 | 0.00862 | 30 | 28 | 3.094 | 49.59 | 0.001343 to 0.05200 | 0.0324 |
| | Post8 | 0.0363 | 0.0303 | 0.005992 | 0.005649 | 30 | 28 | 1.061 | 54.39 | -0.01054 to 0.02252 | >0.9999 |
| **Pert9$_L$** | Young - Older | | | | | | | | | | |
| | Base | 0.05127 | 0.05456 | -0.00329 | 0.003844 | 30 | 28 | 0.856 | 50.37 | -0.01457 to 0.007995 | >0.9999 |
| | Pre | 0.04608 | 0.05559 | -0.009507 | 0.005474 | 30 | 28 | 1.737 | 54.12 | -0.02553 to 0.006512 | 0.8808 |
| | Post1 | -0.04829 | -0.02543 | -0.02285 | 0.01568 | 30 | 28 | 1.458 | 43.13 | -0.06924 to 0.02353 | >0.9999 |
| | Post2 | -0.01154 | 0.01246 | -0.024 | 0.01445 | 30 | 28 | 1.661 | 55.82 | -0.06623 to 0.01823 | >0.9999 |
| | Post3 | 0.01292 | 0.004749 | 0.008169 | 0.01474 | 30 | 28 | 0.554 | 54.68 | -0.03496 to 0.05130 | >0.9999 |
| | Post4 | 0.01281 | 0.01211 | 0.0007005 | 0.01152 | 30 | 28 | 0.06081 | 51.34 | -0.03309 to 0.03449 | >0.9999 |
| | Post5 | 0.03754 | 0.02975 | 0.007787 | 0.009943 | 30 | 28 | 0.7831 | 45.95 | -0.02153 to 0.03711 | >0.9999 |
| | Post6 | 0.04199 | 0.03516 | 0.006832 | 0.008266 | 30 | 28 | 0.8266 | 51.67 | -0.01741 to 0.03107 | >0.9999 |
| | Post7 | 0.05671 | 0.04977 | 0.006938 | 0.007204 | 30 | 28 | 0.963 | 55.17 | -0.01413 to 0.02800 | >0.9999 |
| | Post8 | 0.05212 | 0.04842 | 0.003693 | 0.006066 | 30 | 28 | 0.6088 | 54.88 | -0.01405 to 0.02143 | >0.9999 |

**eTable 3: Bonferroni's multiple comparison test results comparing recovery steps to baseline in Pert1$_R$, Pert2$_L$ and Pert9$_L$ for BoS in the young and older adults**

| Perturbation | Test details | Mean 1 | Mean 2 | Mean Diff. | SE of diff. | N1 | N2 | t | DF | 95.00% CI of diff. | Adjusted P Value |
|---|---|---|---|---|---|---|---|---|---|---|---|
| **Pert1$_R$** | Young | | | | | | | | | | |
| | Base vs. Pre | 0.6682 | 0.6668 | 0.001409 | 0.003597 | 30 | 30 | 0.3916 | 29 | -0.009367 to 0.01218 | >0.9999 |
| | Base vs. Post1 | 0.6682 | 0.7997 | -0.1315 | 0.01834 | 30 | 30 | 7.173 | 29 | -0.1865 to -0.07660 | <0.0001 |
| | Base vs. Post2 | 0.6682 | 0.3485 | 0.3197 | 0.03274 | 30 | 30 | 9.765 | 29 | 0.2216 to 0.4178 | <0.0001 |
| | Base vs. Post3 | 0.6682 | 0.6344 | 0.03379 | 0.02678 | 30 | 30 | 1.261 | 29 | -0.04646 to 0.1140 | >0.9999 |
| | Base vs. Post4 | 0.6682 | 0.7061 | -0.03789 | 0.01115 | 30 | 30 | 3.398 | 29 | -0.07129 to -0.004487 | 0.0179 |
| | Base vs. Post5 | 0.6682 | 0.6424 | 0.02572 | 0.009957 | 30 | 30 | 2.583 | 29 | -0.004110 to 0.05555 | 0.1359 |
| | Base vs. Post6 | 0.6682 | 0.6505 | 0.01764 | 0.008043 | 30 | 30 | 2.194 | 29 | -0.006451 to 0.04174 | 0.3277 |
| | Base vs. Post7 | 0.6682 | 0.6365 | 0.03167 | 0.006189 | 30 | 30 | 5.117 | 29 | 0.01313 to 0.05021 | 0.0002 |
| | Base vs. Post8 | 0.6682 | 0.6511 | 0.01701 | 0.00671 | 30 | 30 | 2.536 | 29 | -0.003088 to 0.03712 | 0.1518 |
| | Older | | | | | | | | | | |
| | Base vs. Pre | 0.6438 | 0.64 | 0.003826 | 0.004034 | 27 | 27 | 0.9484 | 26 | -0.008373 to 0.01603 | >0.9999 |
| | Base vs. Post1 | 0.6438 | 0.7432 | -0.09937 | 0.01022 | 27 | 27 | 9.723 | 26 | -0.1303 to -0.06847 | <0.0001 |
| | Base vs. Post2 | 0.6438 | 0.2436 | 0.4002 | 0.03247 | 27 | 27 | 12.33 | 26 | 0.3020 to 0.4984 | <0.0001 |
| | Base vs. Post3 | 0.6438 | 0.5265 | 0.1173 | 0.02718 | 27 | 27 | 4.318 | 26 | 0.03516 to 0.1995 | 0.0018 |
| | Base vs. Post4 | 0.6438 | 0.6003 | 0.04344 | 0.02796 | 27 | 27 | 1.554 | 26 | -0.04109 to 0.1280 | >0.9999 |
| | Base vs. Post5 | 0.6438 | 0.6205 | 0.0233 | 0.01868 | 27 | 27 | 1.248 | 26 | -0.03317 to 0.07977 | >0.9999 |
| | Base vs. Post6 | 0.6438 | 0.6482 | -0.004443 | 0.01213 | 27 | 27 | 0.3664 | 26 | -0.04111 to 0.03222 | >0.9999 |
| | Base vs. Post7 | 0.6438 | 0.6157 | 0.02805 | 0.01354 | 27 | 27 | 2.071 | 26 | -0.01290 to 0.06899 | 0.4355 |
| | Base vs. Post8 | 0.6438 | 0.629 | 0.01479 | 0.01053 | 27 | 27 | 1.404 | 26 | -0.01705 to 0.04663 | >0.9999 |
| **Pert2$_L$** | Young | | | | | | | | | | |
| | Base vs. Pre | 0.6635 | 0.6556 | 0.007867 | 0.003805 | 30 | 30 | 2.068 | 29 | -0.003531 to 0.01927 | 0.4292 |
| | Base vs. Post1 | 0.6635 | 0.8094 | -0.1459 | 0.01811 | 30 | 30 | 8.055 | 29 | -0.2002 to -0.09164 | <0.0001 |
| | Base vs. Post2 | 0.6635 | 0.4273 | 0.2362 | 0.03137 | 30 | 30 | 7.53 | 29 | 0.1422 to 0.3302 | <0.0001 |
| | Base vs. Post3 | 0.6635 | 0.637 | 0.02653 | 0.01729 | 30 | 30 | 1.534 | 29 | -0.02527 to 0.07833 | >0.9999 |
| | Base vs. Post4 | 0.6635 | 0.7137 | -0.0502 | 0.01132 | 30 | 30 | 4.436 | 29 | -0.08409 to -0.01630 | 0.0011 |
| | Base vs. Post5 | 0.6635 | 0.6667 | -0.003187 | 0.008528 | 30 | 30 | 0.3737 | 29 | -0.02874 to 0.02236 | >0.9999 |
| | Base vs. Post6 | 0.6635 | 0.6543 | 0.009177 | 0.007922 | 30 | 30 | 1.158 | 29 | -0.01456 to 0.03291 | >0.9999 |
| | Base vs. Post7 | 0.6635 | 0.6452 | 0.01832 | 0.008316 | 30 | 30 | 2.203 | 29 | -0.006596 to 0.04323 | 0.3215 |
| | Base vs. Post8 | 0.6635 | 0.6536 | 0.009872 | 0.005144 | 30 | 30 | 1.919 | 29 | -0.005541 to 0.02528 | 0.584 |
| | Older | | | | | | | | | | |
| | Base vs. Pre | 0.6371 | 0.6317 | 0.005399 | 0.005054 | 28 | 28 | 1.068 | 27 | -0.009831 to 0.02063 | >0.9999 |
| | Base vs. Post1 | 0.6371 | 0.7397 | -0.1026 | 0.02012 | 28 | 28 | 5.1 | 27 | -0.1632 to -0.04197 | 0.0002 |
| | Base vs. Post2 | 0.6371 | 0.2684 | 0.3686 | 0.02456 | 28 | 28 | 15.01 | 27 | 0.2946 to 0.4426 | <0.0001 |
| | Base vs. Post3 | 0.6371 | 0.5438 | 0.09323 | 0.01554 | 28 | 28 | 6 | 27 | 0.04640 to 0.1401 | <0.0001 |
| | Base vs. Post4 | 0.6371 | 0.5977 | 0.03941 | 0.02513 | 28 | 28 | 1.568 | 27 | -0.03633 to 0.1152 | >0.9999 |

| | | | | | | | | | | | |
|---|---|---|---|---|---|---|---|---|---|---|---|
| | Base vs. Post5 | 0.6371 | 0.6382 | -0.00111 | 0.01373 | 28 | 28 | 0.0809 | 27 | -0.04248 to 0.04026 | >0.9999 |
| | Base vs. Post6 | 0.6371 | 0.6621 | -0.025 | 0.01198 | 28 | 28 | 2.087 | 27 | -0.06110 to 0.01110 | 0.4183 |
| | Base vs. Post7 | 0.6371 | 0.6355 | 0.001548 | 0.008807 | 28 | 28 | 0.1758 | 27 | -0.02500 to 0.02809 | >0.9999 |
| | Base vs. Post8 | 0.6371 | 0.6335 | 0.00356 | 0.006141 | 28 | 28 | 0.5797 | 27 | -0.01495 to 0.02207 | >0.9999 |
| **Pert9$_L$** | Young | | | | | | | | | | |
| | Base vs. Pre | 0.6725 | 0.6671 | 0.005362 | 0.00302 | 30 | 30 | 1.776 | 29 | -0.003684 to 0.01441 | 0.7762 |
| | Base vs. Post1 | 0.6725 | 0.7637 | -0.09119 | 0.02338 | 30 | 30 | 3.899 | 29 | -0.1612 to -0.02113 | 0.0047 |
| | Base vs. Post2 | 0.6725 | 0.5826 | 0.08989 | 0.01879 | 30 | 30 | 4.785 | 29 | 0.03361 to 0.1462 | 0.0004 |
| | Base vs. Post3 | 0.6725 | 0.6425 | 0.02999 | 0.01391 | 30 | 30 | 2.156 | 29 | -0.01167 to 0.07165 | 0.3553 |
| | Base vs. Post4 | 0.6725 | 0.6992 | -0.02667 | 0.007545 | 30 | 30 | 3.535 | 29 | -0.04927 to -0.004066 | 0.0125 |
| | Base vs. Post5 | 0.6725 | 0.673 | -0.0004707 | 0.00594 | 30 | 30 | 0.0792 | 29 | -0.01827 to 0.01733 | >0.9999 |
| | Base vs. Post6 | 0.6725 | 0.6694 | 0.003134 | 0.004412 | 30 | 30 | 0.7103 | 29 | -0.01009 to 0.01635 | >0.9999 |
| | Base vs. Post7 | 0.6725 | 0.6636 | 0.008934 | 0.004477 | 30 | 30 | 1.996 | 29 | -0.004478 to 0.02235 | 0.4989 |
| | Base vs. Post8 | 0.6725 | 0.6638 | 0.008676 | 0.004403 | 30 | 30 | 1.97 | 29 | -0.004516 to 0.02187 | 0.5258 |
| | Older | | | | | | | | | | |
| | Base vs. Pre | 0.642 | 0.6464 | -0.004371 | 0.004482 | 28 | 28 | 0.9753 | 27 | -0.01788 to 0.009136 | >0.9999 |
| | Base vs. Post1 | 0.642 | 0.7138 | -0.07178 | 0.02028 | 28 | 28 | 3.539 | 27 | -0.1329 to -0.01065 | 0.0133 |
| | Base vs. Post2 | 0.642 | 0.5109 | 0.1311 | 0.026 | 28 | 28 | 5.043 | 27 | 0.05276 to 0.2095 | 0.0002 |
| | Base vs. Post3 | 0.642 | 0.56 | 0.08201 | 0.01619 | 28 | 28 | 5.067 | 27 | 0.03323 to 0.1308 | 0.0002 |
| | Base vs. Post4 | 0.642 | 0.6414 | 0.000607 | 0.01175 | 28 | 28 | 0.0516 | 27 | -0.03481 to 0.03603 | >0.9999 |
| | Base vs. Post5 | 0.642 | 0.6302 | 0.01175 | 0.01323 | 28 | 28 | 0.8883 | 27 | -0.02811 to 0.05161 | >0.9999 |
| | Base vs. Post6 | 0.642 | 0.6591 | -0.01716 | 0.008674 | 28 | 28 | 1.978 | 27 | -0.04330 to 0.008984 | 0.524 |
| | Base vs. Post7 | 0.642 | 0.6342 | 0.007775 | 0.007322 | 28 | 28 | 1.062 | 27 | -0.01429 to 0.02984 | >0.9999 |
| | Base vs. Post8 | 0.642 | 0.6454 | -0.003409 | 0.006732 | 28 | 28 | 0.5064 | 27 | -0.02370 to 0.01688 | >0.9999 |

**eTable 4: Bonferroni's multiple comparison test results comparing the young and older adults in Pert1$_R$, Pert2$_L$ and Pert9$_L$ for BoS**

| Perturbation | Test details | Mean 1 | Mean 2 | Mean Diff. | SE of diff. | N1 | N2 | t | DF | 95.00% CI of diff. | Adjusted P Value |
|---|---|---|---|---|---|---|---|---|---|---|---|
| **Pert1$_R$** | Young - Older | | | | | | | | | | |
| | Base | 0.6682 | 0.6438 | 0.02437 | 0.01305 | 30 | 27 | 1.868 | 43.27 | -0.01423 to 0.06297 | 0.686 |
| | Pre | 0.6668 | 0.64 | 0.02679 | 0.01355 | 30 | 27 | 1.977 | 46.18 | -0.01317 to 0.06674 | 0.5407 |
| | Post1 | 0.7997 | 0.7432 | 0.05654 | 0.02469 | 30 | 27 | 2.29 | 53.97 | -0.01574 to 0.1288 | 0.2598 |
| | Post2 | 0.3485 | 0.2436 | 0.1049 | 0.04659 | 30 | 27 | 2.252 | 54.98 | -0.03136 to 0.2411 | 0.2837 |
| | Post3 | 0.6344 | 0.5265 | 0.1079 | 0.04081 | 30 | 27 | 2.644 | 52.64 | -0.01167 to 0.2275 | 0.1076 |
| | Post4 | 0.7061 | 0.6003 | 0.1057 | 0.03602 | 30 | 27 | 2.935 | 35.84 | -0.002043 to 0.2135 | 0.0579 |
| | Post5 | 0.6424 | 0.6205 | 0.02195 | 0.02719 | 30 | 27 | 0.8073 | 37.35 | -0.05917 to 0.1031 | >0.9999 |
| | Post6 | 0.6505 | 0.6482 | 0.002281 | 0.02124 | 30 | 27 | 0.1074 | 46.4 | -0.06031 to 0.06487 | >0.9999 |
| | Post7 | 0.6365 | 0.6157 | 0.02074 | 0.02275 | 30 | 27 | 0.912 | 37.41 | -0.04711 to 0.08860 | >0.9999 |
| | Post8 | 0.6511 | 0.629 | 0.02214 | 0.01883 | 30 | 27 | 1.176 | 45.9 | -0.03340 to 0.07769 | >0.9999 |
| **Pert2$_L$** | Young - Older | | | | | | | | | | |
| | Base | 0.6635 | 0.6371 | 0.02645 | 0.01339 | 30 | 28 | 1.975 | 44.29 | -0.01312 to 0.06602 | 0.5452 |
| | Pre | 0.6556 | 0.6317 | 0.02398 | 0.01402 | 30 | 28 | 1.711 | 46.85 | -0.01731 to 0.06528 | 0.9367 |
| | Post1 | 0.8094 | 0.7397 | 0.06976 | 0.02983 | 30 | 28 | 2.339 | 54.26 | -0.01753 to 0.1571 | 0.2307 |
| | Post2 | 0.4273 | 0.2684 | 0.1588 | 0.04123 | 30 | 28 | 3.852 | 54.32 | 0.03818 to 0.2795 | 0.0031 |
| | Post3 | 0.637 | 0.5438 | 0.09315 | 0.02674 | 30 | 28 | 3.484 | 50.05 | 0.01464 to 0.1717 | 0.0104 |
| | Post4 | 0.7137 | 0.5977 | 0.1161 | 0.03218 | 30 | 28 | 3.607 | 37.28 | 0.02004 to 0.2121 | 0.009 |
| | Post5 | 0.6667 | 0.6382 | 0.02853 | 0.02237 | 30 | 28 | 1.275 | 46.58 | -0.03739 to 0.09445 | >0.9999 |
| | Post6 | 0.6543 | 0.6621 | -0.007722 | 0.01921 | 30 | 28 | 0.4021 | 47.34 | -0.06428 to 0.04883 | >0.9999 |
| | Post7 | 0.6452 | 0.6355 | 0.009682 | 0.0169 | 30 | 28 | 0.573 | 52.49 | -0.03983 to 0.05920 | >0.9999 |
| | Post8 | 0.6536 | 0.6335 | 0.02014 | 0.01526 | 30 | 28 | 1.319 | 42.98 | -0.02503 to 0.06531 | >0.9999 |
| **Pert9$_L$** | Young - Older | | | | | | | | | | |
| | Base | 0.6725 | 0.642 | 0.03052 | 0.01208 | 30 | 28 | 2.526 | 49.43 | -0.004983 to 0.06601 | 0.1479 |
| | Pre | 0.6671 | 0.6464 | 0.02078 | 0.01344 | 30 | 28 | 1.546 | 47.59 | -0.01879 to 0.06036 | >0.9999 |
| | Post1 | 0.7637 | 0.7138 | 0.04992 | 0.032 | 30 | 28 | 1.56 | 55.5 | -0.04364 to 0.1435 | >0.9999 |
| | Post2 | 0.5826 | 0.5109 | 0.07175 | 0.03598 | 30 | 28 | 1.994 | 49.27 | -0.03401 to 0.1775 | 0.5171 |
| | Post3 | 0.6425 | 0.56 | 0.08253 | 0.02414 | 30 | 28 | 3.419 | 52.12 | 0.01177 to 0.1533 | 0.0123 |
| | Post4 | 0.6992 | 0.6414 | 0.05779 | 0.01735 | 30 | 28 | 3.331 | 54.35 | 0.007018 to 0.1086 | 0.0156 |
| | Post5 | 0.673 | 0.6302 | 0.04274 | 0.01751 | 30 | 28 | 2.44 | 45.44 | -0.008942 to 0.09441 | 0.1865 |
| | Post6 | 0.6694 | 0.6591 | 0.01022 | 0.01615 | 30 | 28 | 0.633 | 47.71 | -0.03732 to 0.05777 | >0.9999 |
| | Post7 | 0.6636 | 0.6342 | 0.02936 | 0.01478 | 30 | 28 | 1.986 | 45.79 | -0.01424 to 0.07296 | 0.5306 |
| | Post8 | 0.6638 | 0.6454 | 0.01843 | 0.01492 | 30 | 28 | 1.236 | 44.75 | -0.02562 to 0.06248 | >0.9999 |

**eTable 5: Bonferroni's multiple comparison test results comparing recovery steps to baseline in Pert1$_R$, Pert2$_L$ and Pert9$_L$ for X$_{CoM}$ in the young and older adults**

| Perturbation | Test details | Mean 1 | Mean 2 | Mean Diff. | SE of diff. | N1 | N2 | t | DF | 95.00% CI of diff. | Adjusted P Value |
|---|---|---|---|---|---|---|---|---|---|---|---|
| **Pert1$_R$** | Young | | | | | | | | | | |
| | Base vs. Pre | 0.6217 | 0.6173 | 0.004363 | 0.005348 | 30 | 30 | 0.8158 | 29 | -0.01166 to 0.02038 | >0.9999 |
| | Base vs. Post1 | 0.6217 | 0.8191 | -0.1974 | 0.01943 | 30 | 30 | 10.16 | 29 | -0.2557 to -0.1392 | <0.0001 |
| | Base vs. Post2 | 0.6217 | 0.3928 | 0.2289 | 0.02759 | 30 | 30 | 8.294 | 29 | 0.1462 to 0.3115 | <0.0001 |
| | Base vs. Post3 | 0.6217 | 0.6951 | -0.07339 | 0.02688 | 30 | 30 | 2.731 | 29 | -0.1539 to 0.007129 | 0.0958 |
| | Base vs. Post4 | 0.6217 | 0.7456 | -0.124 | 0.01765 | 30 | 30 | 7.024 | 29 | -0.1768 to -0.07109 | <0.0001 |
| | Base vs. Post5 | 0.6217 | 0.6405 | -0.01884 | 0.01662 | 30 | 30 | 1.133 | 29 | -0.06864 to 0.03097 | >0.9999 |
| | Base vs. Post6 | 0.6217 | 0.626 | -0.004297 | 0.01361 | 30 | 30 | 0.3157 | 29 | -0.04508 to 0.03649 | >0.9999 |
| | Base vs. Post7 | 0.6217 | 0.6061 | 0.01557 | 0.00955 | 30 | 30 | 1.631 | 29 | -0.01304 to 0.04418 | >0.9999 |
| | Base vs. Post8 | 0.6217 | 0.6114 | 0.01028 | 0.01082 | 30 | 30 | 0.9504 | 29 | -0.02213 to 0.04269 | >0.9999 |
| | Older | | | | | | | | | | |
| | Base vs. Pre | 0.5942 | 0.5942 | -9.535E-06 | 0.005839 | 27 | 27 | 0.0016 | 26 | -0.01767 to 0.01765 | >0.9999 |
| | Base vs. Post1 | 0.5942 | 0.6314 | -0.03724 | 0.01465 | 27 | 27 | 2.542 | 26 | -0.08154 to 0.007056 | 0.1559 |
| | Base vs. Post2 | 0.5942 | 0.2209 | 0.3733 | 0.03143 | 27 | 27 | 11.88 | 26 | 0.2783 to 0.4683 | <0.0001 |
| | Base vs. Post3 | 0.5942 | 0.5272 | 0.06696 | 0.02828 | 27 | 27 | 2.368 | 26 | -0.01856 to 0.1525 | 0.2307 |
| | Base vs. Post4 | 0.5942 | 0.6282 | -0.03402 | 0.0273 | 27 | 27 | 1.246 | 26 | -0.1166 to 0.04853 | >0.9999 |
| | Base vs. Post5 | 0.5942 | 0.6302 | -0.03604 | 0.02071 | 27 | 27 | 1.741 | 26 | -0.09866 to 0.02657 | 0.8421 |
| | Base vs. Post6 | 0.5942 | 0.644 | -0.04984 | 0.0135 | 27 | 27 | 3.692 | 26 | -0.09065 to -0.009022 | 0.0093 |
| | Base vs. Post7 | 0.5942 | 0.6001 | -0.0059 | 0.01252 | 27 | 27 | 0.4712 | 26 | -0.04376 to 0.03196 | >0.9999 |
| | Base vs. Post8 | 0.5942 | 0.6029 | -0.00868 | 0.01323 | 27 | 27 | 0.656 | 26 | -0.04869 to 0.03133 | >0.9999 |
| **Pert2$_L$** | Young | | | | | | | | | | |
| | Base vs. Pre | 0.6201 | 0.6159 | 0.004229 | 0.00424 | 30 | 30 | 0.9974 | 29 | -0.008474 to 0.01693 | >0.9999 |
| | Base vs. Post1 | 0.6201 | 0.8288 | -0.2087 | 0.01824 | 30 | 30 | 11.44 | 29 | -0.2634 to -0.1541 | <0.0001 |
| | Base vs. Post2 | 0.6201 | 0.4474 | 0.1727 | 0.02634 | 30 | 30 | 6.556 | 29 | 0.09378 to 0.2516 | <0.0001 |
| | Base vs. Post3 | 0.6201 | 0.6679 | -0.04785 | 0.02408 | 30 | 30 | 1.987 | 29 | -0.1200 to 0.02430 | 0.508 |
| | Base vs. Post4 | 0.6201 | 0.7533 | -0.1333 | 0.01764 | 30 | 30 | 7.553 | 29 | -0.1861 to -0.08040 | <0.0001 |
| | Base vs. Post5 | 0.6201 | 0.6604 | -0.04032 | 0.01577 | 30 | 30 | 2.558 | 29 | -0.08755 to 0.006911 | 0.1443 |
| | Base vs. Post6 | 0.6201 | 0.6277 | -0.007645 | 0.01022 | 30 | 30 | 0.7478 | 29 | -0.03827 to 0.02298 | >0.9999 |
| | Base vs. Post7 | 0.6201 | 0.6007 | 0.0194 | 0.009114 | 30 | 30 | 2.128 | 29 | -0.007906 to 0.04670 | 0.3773 |
| | Base vs. Post8 | 0.6201 | 0.6173 | 0.002745 | 0.007312 | 30 | 30 | 0.3754 | 29 | -0.01916 to 0.02465 | >0.9999 |
| | Older | | | | | | | | | | |
| | Base vs. Pre | 0.5893 | 0.5877 | 0.001586 | 0.006807 | 28 | 28 | 0.233 | 27 | -0.01893 to 0.02210 | >0.9999 |
| | Base vs. Post1 | 0.5893 | 0.672 | -0.08275 | 0.02284 | 28 | 28 | 3.623 | 27 | -0.1516 to -0.01392 | 0.0107 |
| | Base vs. Post2 | 0.5893 | 0.2663 | 0.3229 | 0.0189 | 28 | 28 | 17.08 | 27 | 0.2660 to 0.3799 | <0.0001 |
| | Base vs. Post3 | 0.5893 | 0.5299 | 0.05935 | 0.01948 | 28 | 28 | 3.046 | 27 | 0.0006364 to 0.1181 | 0.0461 |
| | Base vs. Post4 | 0.5893 | 0.6351 | -0.04588 | 0.02789 | 28 | 28 | 1.645 | 27 | -0.1299 to 0.03816 | >0.9999 |

| | | | | | | | | | | | |
|---|---|---|---|---|---|---|---|---|---|---|---|
| | Base vs. Post5 | 0.5893 | 0.637 | -0.04778 | 0.01887 | 28 | 28 | 2.533 | 27 | -0.1046 to 0.009073 | 0.1569 |
| | Base vs. Post6 | 0.5893 | 0.6674 | -0.07809 | 0.01426 | 28 | 28 | 5.475 | 27 | -0.1211 to -0.03511 | <0.0001 |
| | Base vs. Post7 | 0.5893 | 0.6177 | -0.02842 | 0.01173 | 28 | 28 | 2.423 | 27 | -0.06376 to 0.006925 | 0.2013 |
| | Base vs. Post8 | 0.5893 | 0.6032 | -0.01394 | 0.00787 | 28 | 28 | 1.771 | 27 | -0.03765 to 0.009781 | 0.7908 |
| **Pert9$_L$** | Young | | | | | | | | | | |
| | Base vs. Pre | 0.6212 | 0.6211 | 0.0001787 | 0.005884 | 30 | 30 | 0.0304 | 29 | -0.01745 to 0.01781 | >0.9999 |
| | Base vs. Post1 | 0.6212 | 0.812 | -0.1907 | 0.02334 | 30 | 30 | 8.171 | 29 | -0.2607 to -0.1208 | <0.0001 |
| | Base vs. Post2 | 0.6212 | 0.5942 | 0.02708 | 0.01529 | 30 | 30 | 1.771 | 29 | -0.01873 to 0.07289 | 0.784 |
| | Base vs. Post3 | 0.6212 | 0.6296 | -0.008359 | 0.01927 | 30 | 30 | 0.4337 | 29 | -0.06610 to 0.04938 | >0.9999 |
| | Base vs. Post4 | 0.6212 | 0.6864 | -0.06513 | 0.01207 | 30 | 30 | 5.397 | 29 | -0.1013 to -0.02897 | <0.0001 |
| | Base vs. Post5 | 0.6212 | 0.6354 | -0.0142 | 0.009939 | 30 | 30 | 1.429 | 29 | -0.04398 to 0.01557 | >0.9999 |
| | Base vs. Post6 | 0.6212 | 0.6274 | -0.006144 | 0.007752 | 30 | 30 | 0.7926 | 29 | -0.02937 to 0.01708 | >0.9999 |
| | Base vs. Post7 | 0.6212 | 0.6069 | 0.01438 | 0.007794 | 30 | 30 | 1.845 | 29 | -0.008973 to 0.03772 | 0.678 |
| | Base vs. Post8 | 0.6212 | 0.6117 | 0.009525 | 0.007469 | 30 | 30 | 1.275 | 29 | -0.01285 to 0.03190 | >0.9999 |
| | Older | | | | | | | | | | |
| | Base vs. Pre | 0.5874 | 0.5908 | -0.003338 | 0.006542 | 28 | 28 | 0.5102 | 27 | -0.02305 to 0.01638 | >0.9999 |
| | Base vs. Post1 | 0.5874 | 0.7392 | -0.1518 | 0.02003 | 28 | 28 | 7.578 | 27 | -0.2121 to -0.09141 | <0.0001 |
| | Base vs. Post2 | 0.5874 | 0.4984 | 0.08902 | 0.02295 | 28 | 28 | 3.879 | 27 | 0.01986 to 0.1582 | 0.0055 |
| | Base vs. Post3 | 0.5874 | 0.5552 | 0.0322 | 0.01658 | 28 | 28 | 1.942 | 27 | -0.01778 to 0.08217 | 0.5641 |
| | Base vs. Post4 | 0.5874 | 0.6293 | -0.04184 | 0.01663 | 28 | 28 | 2.517 | 27 | -0.09195 to 0.008264 | 0.1628 |
| | Base vs. Post5 | 0.5874 | 0.6005 | -0.01306 | 0.01649 | 28 | 28 | 0.792 | 27 | -0.06276 to 0.03664 | >0.9999 |
| | Base vs. Post6 | 0.5874 | 0.624 | -0.03656 | 0.01415 | 28 | 28 | 2.583 | 27 | -0.07921 to 0.006095 | 0.1398 |
| | Base vs. Post7 | 0.5874 | 0.5844 | 0.002988 | 0.009635 | 28 | 28 | 0.3102 | 27 | -0.02605 to 0.03203 | >0.9999 |
| | Base vs. Post8 | 0.5874 | 0.597 | -0.009543 | 0.009634 | 28 | 28 | 0.9906 | 27 | -0.03858 to 0.01949 | >0.9999 |

**eTable 6: Bonferroni's multiple comparison test results comparing the young and older adults in Pert1$_R$, Pert2$_L$ and Pert9$_L$ for X$_{CoM}$**

| Perturbation | Test details | Mean 1 | Mean 2 | Mean Diff. | SE of diff. | N1 | N2 | t | DF | 95.00% CI of diff. | Adjusted P Value |
|---|---|---|---|---|---|---|---|---|---|---|---|
| **Pert1$_R$** | Young - Older | | | | | | | | | | |
| | Base | 0.6217 | 0.5942 | 0.02748 | 0.01341 | 30 | 27 | 2.049 | 40.77 | -0.01232 to 0.06728 | 0.4692 |
| | Pre | 0.6173 | 0.5942 | 0.02311 | 0.01418 | 30 | 27 | 1.629 | 47 | -0.01867 to 0.06488 | >0.9999 |
| | Post1 | 0.8191 | 0.6314 | 0.1877 | 0.02587 | 30 | 27 | 7.255 | 53.23 | 0.1119 to 0.2634 | <0.0001 |
| | Post2 | 0.3928 | 0.2209 | 0.1719 | 0.04256 | 30 | 27 | 4.04 | 52.7 | 0.04722 to 0.2966 | 0.0017 |
| | Post3 | 0.6951 | 0.5272 | 0.1678 | 0.04267 | 30 | 27 | 3.933 | 52.05 | 0.04271 to 0.2929 | 0.0025 |
| | Post4 | 0.7456 | 0.6282 | 0.1174 | 0.03759 | 30 | 27 | 3.123 | 44.95 | 0.006426 to 0.2284 | 0.0313 |
| | Post5 | 0.6405 | 0.6302 | 0.01027 | 0.03197 | 30 | 27 | 0.3213 | 47.22 | -0.08387 to 0.1044 | >0.9999 |
| | Post6 | 0.626 | 0.644 | -0.01806 | 0.02611 | 30 | 27 | 0.6916 | 52.83 | -0.09457 to 0.05845 | >0.9999 |
| | Post7 | 0.6061 | 0.6001 | 0.006003 | 0.02362 | 30 | 27 | 0.2541 | 44.82 | -0.06374 to 0.07575 | >0.9999 |
| | Post8 | 0.6114 | 0.6029 | 0.008517 | 0.02263 | 30 | 27 | 0.3763 | 51.81 | -0.05784 to 0.07487 | >0.9999 |
| **Pert2$_L$** | Young - Older | | | | | | | | | | |
| | Base | 0.6201 | 0.5893 | 0.03083 | 0.01398 | 30 | 28 | 2.205 | 43.42 | -0.01052 to 0.07218 | 0.328 |
| | Pre | 0.6159 | 0.5877 | 0.02819 | 0.01531 | 30 | 28 | 1.841 | 46.81 | -0.01692 to 0.07329 | 0.7196 |
| | Post1 | 0.8288 | 0.672 | 0.1568 | 0.0318 | 30 | 28 | 4.93 | 52.57 | 0.06360 to 0.2500 | <0.0001 |
| | Post2 | 0.4474 | 0.2663 | 0.1811 | 0.03541 | 30 | 28 | 5.112 | 54.18 | 0.07741 to 0.2847 | <0.0001 |
| | Post3 | 0.6679 | 0.5299 | 0.138 | 0.03578 | 30 | 28 | 3.858 | 54.81 | 0.03337 to 0.2427 | 0.003 |
| | Post4 | 0.7533 | 0.6351 | 0.1182 | 0.03644 | 30 | 28 | 3.244 | 47.29 | 0.01090 to 0.2255 | 0.0217 |
| | Post5 | 0.6604 | 0.637 | 0.02337 | 0.02945 | 30 | 28 | 0.7933 | 54.4 | -0.06282 to 0.1096 | >0.9999 |
| | Post6 | 0.6277 | 0.6674 | -0.03962 | 0.02214 | 30 | 28 | 1.79 | 51.7 | -0.1045 to 0.02530 | 0.7934 |
| | Post7 | 0.6007 | 0.6177 | -0.01699 | 0.01807 | 30 | 28 | 0.9402 | 54.36 | -0.06986 to 0.03588 | >0.9999 |
| | Post8 | 0.6173 | 0.6032 | 0.01415 | 0.01601 | 30 | 28 | 0.8835 | 46.49 | -0.03304 to 0.06134 | >0.9999 |
| **Pert9$_L$** | Young - Older | | | | | | | | | | |
| | Base | 0.6212 | 0.5874 | 0.03381 | 0.01298 | 30 | 28 | 2.605 | 43.69 | -0.004570 to 0.07218 | 0.1253 |
| | Pre | 0.6211 | 0.5908 | 0.03029 | 0.01509 | 30 | 28 | 2.007 | 48.84 | -0.01408 to 0.07466 | 0.5028 |
| | Post1 | 0.812 | 0.7392 | 0.07278 | 0.03309 | 30 | 28 | 2.199 | 55.97 | -0.02394 to 0.1695 | 0.3202 |
| | Post2 | 0.5942 | 0.4984 | 0.09575 | 0.03315 | 30 | 28 | 2.888 | 45.77 | -0.002033 to 0.1935 | 0.059 |
| | Post3 | 0.6296 | 0.5552 | 0.07436 | 0.02908 | 30 | 28 | 2.557 | 55.99 | -0.01061 to 0.1593 | 0.1328 |
| | Post4 | 0.6864 | 0.6293 | 0.05709 | 0.02222 | 30 | 28 | 2.57 | 54.15 | -0.007934 to 0.1221 | 0.1297 |
| | Post5 | 0.6354 | 0.6005 | 0.03495 | 0.02058 | 30 | 28 | 1.698 | 49.95 | -0.02549 to 0.09538 | 0.9564 |
| | Post6 | 0.6274 | 0.624 | 0.003393 | 0.02126 | 30 | 28 | 0.1595 | 46.66 | -0.05927 to 0.06605 | >0.9999 |
| | Post7 | 0.6069 | 0.5844 | 0.02242 | 0.0179 | 30 | 28 | 1.252 | 50.23 | -0.03014 to 0.07498 | >0.9999 |
| | Post8 | 0.6117 | 0.597 | 0.01474 | 0.01775 | 30 | 28 | 0.8302 | 43.64 | -0.03776 to 0.06723 | >0.9999 |

**eTable 7: Bonferroni's multiple comparison test results comparing recovery steps to baseline in Pert1$_R$, Pert2$_L$ and Pert9$_L$ for MoS$_{ML}$ in the young and older adults**

| Perturbation | Test details | Mean 1 | Mean 2 | Mean Diff. | SE of diff. | N1 | N2 | t | DF | 95.00% CI of diff. | Adjusted P Value |
|---|---|---|---|---|---|---|---|---|---|---|---|
| **Pert1$_R$** | Young | | | | | | | | | | |
| | Base vs. Pre | 0.00993 | 0.01381 | -0.003881 | 0.002857 | 30 | 30 | 1.358 | 29 | -0.01244 to 0.004678 | >0.9999 |
| | Base vs. Post1 | 0.00993 | -0.01824 | 0.02817 | 0.006462 | 30 | 30 | 4.359 | 29 | 0.008809 to 0.04753 | 0.0013 |
| | Base vs. Post2 | 0.00993 | 0.05714 | -0.04721 | 0.007052 | 30 | 30 | 6.694 | 29 | -0.06834 to -0.02608 | <0.0001 |
| | Base vs. Post3 | 0.00993 | 0.02722 | -0.01729 | 0.006142 | 30 | 30 | 2.815 | 29 | -0.03569 to 0.001108 | 0.078 |
| | Base vs. Post4 | 0.00993 | 0.01635 | -0.006418 | 0.004151 | 30 | 30 | 1.546 | 29 | -0.01885 to 0.006019 | >0.9999 |
| | Base vs. Post5 | 0.00993 | 0.01654 | -0.006613 | 0.003173 | 30 | 30 | 2.084 | 29 | -0.01612 to 0.002892 | 0.4144 |
| | Base vs. Post6 | 0.00993 | 0.01908 | -0.009145 | 0.005265 | 30 | 30 | 1.737 | 29 | -0.02492 to 0.006629 | 0.8372 |
| | Base vs. Post7 | 0.00993 | 0.01768 | -0.007747 | 0.002813 | 30 | 30 | 2.754 | 29 | -0.01617 to 0.0006799 | 0.0905 |
| | Base vs. Post8 | 0.00993 | 0.02129 | -0.01136 | 0.003613 | 30 | 30 | 3.145 | 29 | -0.02219 to -0.0005390 | 0.0343 |
| | Older | | | | | | | | | | |
| | Base vs. Pre | 0.01644 | 0.02095 | -0.004507 | 0.00331 | 27 | 27 | 1.362 | 26 | -0.01452 to 0.005503 | >0.9999 |
| | Base vs. Post1 | 0.01644 | 0.005855 | 0.01058 | 0.008328 | 27 | 27 | 1.271 | 26 | -0.01460 to 0.03576 | >0.9999 |
| | Base vs. Post2 | 0.01644 | 0.0454 | -0.02897 | 0.00871 | 27 | 27 | 3.325 | 26 | -0.05530 to -0.002628 | 0.0237 |
| | Base vs. Post3 | 0.01644 | 0.03473 | -0.01829 | 0.009911 | 27 | 27 | 1.845 | 26 | -0.04826 to 0.01168 | 0.6876 |
| | Base vs. Post4 | 0.01644 | 0.04115 | -0.02471 | 0.005466 | 27 | 27 | 4.521 | 26 | -0.04124 to -0.008185 | 0.0011 |
| | Base vs. Post5 | 0.01644 | 0.02488 | -0.00844 | 0.00469 | 27 | 27 | 1.799 | 26 | -0.02262 to 0.005742 | 0.7521 |
| | Base vs. Post6 | 0.01644 | 0.03278 | -0.01634 | 0.004258 | 27 | 27 | 3.837 | 26 | -0.02922 to -0.003464 | 0.0064 |
| | Base vs. Post7 | 0.01644 | 0.02323 | -0.006792 | 0.004249 | 27 | 27 | 1.598 | 26 | -0.01964 to 0.006056 | >0.9999 |
| | Base vs. Post8 | 0.01644 | 0.02829 | -0.01185 | 0.004343 | 27 | 27 | 2.729 | 26 | -0.02499 to 0.001278 | 0.1011 |
| **Pert2$_L$** | Young | | | | | | | | | | |
| | Base vs. Pre | 0.01463 | 0.01294 | 0.001691 | 0.00324 | 30 | 30 | 0.5218 | 29 | -0.008015 to 0.01140 | >0.9999 |
| | Base vs. Post1 | 0.01463 | -0.01619 | 0.03082 | 0.009414 | 30 | 30 | 3.274 | 29 | 0.002616 to 0.05902 | 0.0247 |
| | Base vs. Post2 | 0.01463 | 0.06054 | -0.0459 | 0.006201 | 30 | 30 | 7.402 | 29 | -0.06448 to -0.02732 | <0.0001 |
| | Base vs. Post3 | 0.01463 | 0.03341 | -0.01877 | 0.004804 | 30 | 30 | 3.908 | 29 | -0.03317 to -0.004380 | 0.0046 |
| | Base vs. Post4 | 0.01463 | 0.0199 | -0.00527 | 0.003388 | 30 | 30 | 1.555 | 29 | -0.01542 to 0.004880 | >0.9999 |
| | Base vs. Post5 | 0.01463 | 0.01923 | -0.004594 | 0.003308 | 30 | 30 | 1.389 | 29 | -0.01450 to 0.005316 | >0.9999 |
| | Base vs. Post6 | 0.01463 | 0.02158 | -0.006944 | 0.003387 | 30 | 30 | 2.05 | 29 | -0.01709 to 0.003203 | 0.4453 |
| | Base vs. Post7 | 0.01463 | 0.02254 | -0.007906 | 0.002615 | 30 | 30 | 3.024 | 29 | -0.01574 to -7.218e-005 | 0.0467 |
| | Base vs. Post8 | 0.01463 | 0.01893 | -0.004295 | 0.002056 | 30 | 30 | 2.089 | 29 | -0.01045 to 0.001864 | 0.4104 |
| | Older | | | | | | | | | | |
| | Base vs. Pre | 0.01851 | 0.01986 | -0.00135 | 0.002821 | 28 | 28 | 0.4788 | 27 | -0.009851 to 0.007150 | >0.9999 |
| | Base vs. Post1 | 0.01851 | 0.003911 | 0.0146 | 0.00723 | 28 | 28 | 2.019 | 27 | -0.007189 to 0.03639 | 0.4813 |
| | Base vs. Post2 | 0.01851 | 0.04397 | -0.02545 | 0.00709 | 28 | 28 | 3.59 | 27 | -0.04682 to -0.004086 | 0.0117 |
| | Base vs. Post3 | 0.01851 | 0.04124 | -0.02273 | 0.005476 | 28 | 28 | 4.15 | 27 | -0.03923 to -0.006223 | 0.0027 |
| | Base vs. Post4 | 0.01851 | 0.02997 | -0.01146 | 0.005122 | 28 | 28 | 2.237 | 27 | -0.02690 to 0.003977 | 0.3035 |

| | | | | | | | | | | | |
|---|---|---|---|---|---|---|---|---|---|---|---|
| | Base vs. Post5 | 0.01851 | 0.03564 | -0.01712 | 0.005769 | 28 | 28 | 2.969 | 27 | -0.03451 to 0.0002602 | 0.0558 |
| | Base vs. Post6 | 0.01851 | 0.02377 | -0.005261 | 0.004111 | 28 | 28 | 1.28 | 27 | -0.01765 to 0.007128 | >0.9999 |
| | Base vs. Post7 | 0.01851 | 0.02978 | -0.01126 | 0.004092 | 28 | 28 | 2.753 | 27 | -0.02360 to 0.001068 | 0.0939 |
| | Base vs. Post8 | 0.01851 | 0.02537 | -0.006858 | 0.003129 | 28 | 28 | 2.192 | 27 | -0.01629 to 0.002572 | 0.3348 |
| **Pert9$_L$** | Young | | | | | | | | | | |
| | Base vs. Pre | 0.01167 | 0.01541 | -0.003741 | 0.002662 | 30 | 30 | 1.406 | 29 | -0.01172 to 0.004233 | >0.9999 |
| | Base vs. Post1 | 0.01167 | 0.01429 | -0.002621 | 0.006735 | 30 | 30 | 0.3891 | 29 | -0.02280 to 0.01756 | >0.9999 |
| | Base vs. Post2 | 0.01167 | 0.04776 | -0.03609 | 0.005783 | 30 | 30 | 6.242 | 29 | -0.05342 to -0.01877 | <0.0001 |
| | Base vs. Post3 | 0.01167 | 0.02938 | -0.01772 | 0.004635 | 30 | 30 | 3.822 | 29 | -0.03160 to -0.003830 | 0.0058 |
| | Base vs. Post4 | 0.01167 | 0.01602 | -0.004356 | 0.003774 | 30 | 30 | 1.154 | 29 | -0.01566 to 0.006952 | >0.9999 |
| | Base vs. Post5 | 0.01167 | 0.01403 | -0.002361 | 0.003278 | 30 | 30 | 0.7201 | 29 | -0.01218 to 0.007460 | >0.9999 |
| | Base vs. Post6 | 0.01167 | 0.008163 | 0.003503 | 0.002902 | 30 | 30 | 1.207 | 29 | -0.005190 to 0.01220 | >0.9999 |
| | Base vs. Post7 | 0.01167 | 0.01456 | -0.002889 | 0.003158 | 30 | 30 | 0.9149 | 29 | -0.01235 to 0.006571 | >0.9999 |
| | Base vs. Post8 | 0.01167 | 0.009144 | 0.002523 | 0.002191 | 30 | 30 | 1.151 | 29 | -0.004042 to 0.009087 | >0.9999 |
| | Older | | | | | | | | | | |
| | Base vs. Pre | 0.02119 | 0.01881 | 0.002382 | 0.003569 | 28 | 28 | 0.6672 | 27 | -0.008375 to 0.01314 | >0.9999 |
| | Base vs. Post1 | 0.02119 | -0.001606 | 0.0228 | 0.008087 | 28 | 28 | 2.819 | 27 | -0.001575 to 0.04717 | 0.0802 |
| | Base vs. Post2 | 0.02119 | 0.04653 | -0.02534 | 0.005958 | 28 | 28 | 4.253 | 27 | -0.04330 to -0.007386 | 0.002 |
| | Base vs. Post3 | 0.02119 | 0.04963 | -0.02844 | 0.00501 | 28 | 28 | 5.677 | 27 | -0.04354 to -0.01334 | <0.0001 |
| | Base vs. Post4 | 0.02119 | 0.0268 | -0.005611 | 0.005691 | 28 | 28 | 0.986 | 27 | -0.02276 to 0.01154 | >0.9999 |
| | Base vs. Post5 | 0.02119 | 0.03289 | -0.01169 | 0.003262 | 28 | 28 | 3.585 | 27 | -0.02152 to -0.001864 | 0.0118 |
| | Base vs. Post6 | 0.02119 | 0.02511 | -0.00392 | 0.00281 | 28 | 28 | 1.395 | 27 | -0.01239 to 0.004547 | >0.9999 |
| | Base vs. Post7 | 0.02119 | 0.02618 | -0.004989 | 0.003031 | 28 | 28 | 1.646 | 27 | -0.01412 to 0.004146 | >0.9999 |
| | Base vs. Post8 | 0.02119 | 0.01847 | 0.002723 | 0.00302 | 28 | 28 | 0.9016 | 27 | -0.006379 to 0.01183 | >0.9999 |

**eTable 8: Bonferroni's multiple comparison test results comparing the young and older adults in Pert1$_R$, Pert2$_L$ and Pert9$_L$ for MoS$_{ML}$**

| Perturbation | Test details | Mean 1 | Mean 2 | Mean Diff. | SE of diff. | N1 | N2 | t | DF | 95.00% CI of diff. | Adjusted P Value |
|---|---|---|---|---|---|---|---|---|---|---|---|
| **Pert1$_R$** | Young - Older | | | | | | | | | | |
| | Base | 0.00993 | 0.01644 | -0.006509 | 0.003998 | 30 | 27 | 1.628 | 52.73 | -0.01822 to 0.005206 | >0.9999 |
| | Pre | 0.01381 | 0.02095 | -0.007135 | 0.005738 | 30 | 27 | 1.243 | 53.19 | -0.02394 to 0.009672 | >0.9999 |
| | Post1 | -0.01824 | 0.005855 | -0.02409 | 0.009986 | 30 | 27 | 2.413 | 51.33 | -0.05339 to 0.005199 | 0.1944 |
| | Post2 | 0.05714 | 0.0454 | 0.01174 | 0.01095 | 30 | 27 | 1.072 | 49.5 | -0.02044 to 0.04391 | >0.9999 |
| | Post3 | 0.02722 | 0.03473 | -0.007506 | 0.01242 | 30 | 27 | 0.6042 | 40.22 | -0.04440 to 0.02939 | >0.9999 |
| | Post4 | 0.01635 | 0.04115 | -0.0248 | 0.007797 | 30 | 27 | 3.181 | 50.04 | -0.04770 to -0.001907 | 0.0252 |
| | Post5 | 0.01654 | 0.02488 | -0.008335 | 0.006806 | 30 | 27 | 1.225 | 52.35 | -0.02828 to 0.01161 | >0.9999 |
| | Post6 | 0.01908 | 0.03278 | -0.0137 | 0.008015 | 30 | 27 | 1.71 | 53.68 | -0.03717 to 0.009762 | 0.9308 |
| | Post7 | 0.01768 | 0.02323 | -0.005553 | 0.006228 | 30 | 27 | 0.8916 | 54.22 | -0.02378 to 0.01267 | >0.9999 |
| | Post8 | 0.02129 | 0.02829 | -0.007001 | 0.006483 | 30 | 27 | 1.08 | 51.49 | -0.02602 to 0.01202 | >0.9999 |
| **Pert2$_L$** | Young - Older | | | | | | | | | | |
| | Base | 0.01463 | 0.01851 | -0.003877 | 0.004182 | 30 | 28 | 0.9271 | 55.37 | -0.01610 to 0.008350 | >0.9999 |
| | Pre | 0.01294 | 0.01986 | -0.006918 | 0.005662 | 30 | 28 | 1.222 | 55.99 | -0.02346 to 0.009629 | >0.9999 |
| | Post1 | -0.01619 | 0.003911 | -0.0201 | 0.01263 | 30 | 28 | 1.591 | 50.78 | -0.05717 to 0.01698 | >0.9999 |
| | Post2 | 0.06054 | 0.04397 | 0.01657 | 0.01045 | 30 | 28 | 1.586 | 53.62 | -0.01402 to 0.04716 | >0.9999 |
| | Post3 | 0.03341 | 0.04124 | -0.007828 | 0.007677 | 30 | 28 | 1.02 | 48.24 | -0.03041 to 0.01476 | >0.9999 |
| | Post4 | 0.0199 | 0.02997 | -0.01007 | 0.006901 | 30 | 28 | 1.459 | 50.48 | -0.03033 to 0.01019 | >0.9999 |
| | Post5 | 0.01923 | 0.03564 | -0.01641 | 0.00749 | 30 | 28 | 2.19 | 42.79 | -0.03858 to 0.005764 | 0.3399 |
| | Post6 | 0.02158 | 0.02377 | -0.002194 | 0.006056 | 30 | 28 | 0.3623 | 54.83 | -0.01991 to 0.01552 | >0.9999 |
| | Post7 | 0.02254 | 0.02978 | -0.007235 | 0.006073 | 30 | 28 | 1.191 | 47.56 | -0.02511 to 0.01064 | >0.9999 |
| | Post8 | 0.01893 | 0.02537 | -0.00644 | 0.005667 | 30 | 28 | 1.136 | 53.54 | -0.02303 to 0.01015 | >0.9999 |
| **Pert9$_L$** | Young - Older | | | | | | | | | | |
| | Base | 0.01167 | 0.02119 | -0.009525 | 0.0047 | 30 | 28 | 2.027 | 53.32 | -0.02329 to 0.004239 | 0.4772 |
| | Pre | 0.01541 | 0.01881 | -0.003402 | 0.005727 | 30 | 28 | 0.594 | 55.22 | -0.02015 to 0.01335 | >0.9999 |
| | Post1 | 0.01429 | -0.001606 | 0.01589 | 0.01056 | 30 | 28 | 1.506 | 55.98 | -0.01496 to 0.04674 | >0.9999 |
| | Post2 | 0.04776 | 0.04653 | 0.001227 | 0.00871 | 30 | 28 | 0.1409 | 53.5 | -0.02428 to 0.02673 | >0.9999 |
| | Post3 | 0.02938 | 0.04963 | -0.02025 | 0.006971 | 30 | 28 | 2.905 | 55.99 | -0.04062 to 0.0001226 | 0.0525 |
| | Post4 | 0.01602 | 0.0268 | -0.01078 | 0.007162 | 30 | 28 | 1.505 | 42.74 | -0.03198 to 0.01042 | >0.9999 |
| | Post5 | 0.01403 | 0.03289 | -0.01886 | 0.005902 | 30 | 28 | 3.195 | 52.51 | -0.03615 to -0.001563 | 0.0236 |
| | Post6 | 0.00816 | 0.02511 | -0.01695 | 0.005176 | 30 | 28 | 3.275 | 54.59 | -0.03209 to -0.001806 | 0.0184 |
| | Post7 | 0.01456 | 0.02618 | -0.01162 | 0.005771 | 30 | 28 | 2.014 | 52.19 | -0.02854 to 0.005293 | 0.4915 |
| | Post8 | 0.00914 | 0.01847 | -0.009325 | 0.005245 | 30 | 28 | 1.778 | 46.84 | -0.02478 to 0.006128 | 0.8194 |

**eTable 9: Bonferroni's multiple comparison test results comparing recovery steps to baseline in Pert1$_R$ and Pert10$_R$ for MoS$_{AP}$ in the older adults**

| Test details | Mean 1 | Mean 2 | Mean Diff. | SE of diff. | N1 | N2 | t | DF | 95.00% CI of diff. | Adjusted P Value |
|---|---|---|---|---|---|---|---|---|---|---|
| Pert1$_R$ | | | | | | | | | | |
| Base vs. Pre | 0.0496 | 0.04576 | 0.003836 | 0.003399 | 27 | 27 | 1.128 | 26 | -0.006443 to 0.01411 | >0.9999 |
| Base vs. Post1 | 0.0496 | 0.1117 | -0.06213 | 0.01244 | 27 | 27 | 4.996 | 26 | -0.09973 to -0.02452 | 0.0003 |
| Base vs. Post2 | 0.0496 | 0.02267 | 0.02693 | 0.01389 | 27 | 27 | 1.939 | 26 | -0.01506 to 0.06892 | 0.5709 |
| Base vs. Post3 | 0.0496 | -0.0007875 | 0.05038 | 0.01063 | 27 | 27 | 4.739 | 26 | 0.01824 to 0.08253 | 0.0006 |
| Base vs. Post4 | 0.0496 | -0.02787 | 0.07746 | 0.01084 | 27 | 27 | 7.143 | 26 | 0.04467 to 0.1103 | <0.0001 |
| Base vs. Post5 | 0.0496 | -0.009746 | 0.05934 | 0.01015 | 27 | 27 | 5.846 | 26 | 0.02865 to 0.09004 | <0.0001 |
| Base vs. Post6 | 0.0496 | 0.004204 | 0.04539 | 0.007151 | 27 | 27 | 6.348 | 26 | 0.02377 to 0.06702 | <0.0001 |
| Base vs. Post7 | 0.0496 | 0.01565 | 0.03395 | 0.007473 | 27 | 27 | 4.542 | 26 | 0.01135 to 0.05654 | 0.001 |
| Base vs. Post8 | 0.0496 | 0.02613 | 0.02347 | 0.0077 | 27 | 27 | 3.048 | 26 | 0.0001856 to 0.04675 | 0.0471 |
| Pert10$_R$ | | | | | | | | | | |
| Base vs. Pre | 0.04955 | 0.04616 | 0.003387 | 0.004206 | 27 | 27 | 0.8053 | 26 | -0.009330 to 0.01610 | >0.9999 |
| Base vs. Post1 | 0.04955 | 0.01955 | 0.02999 | 0.01472 | 27 | 27 | 2.038 | 26 | -0.01452 to 0.07450 | 0.467 |
| Base vs. Post2 | 0.04955 | -0.03525 | 0.08479 | 0.01148 | 27 | 27 | 7.388 | 26 | 0.05009 to 0.1195 | <0.0001 |
| Base vs. Post3 | 0.04955 | -0.0292 | 0.07875 | 0.01147 | 27 | 27 | 6.868 | 26 | 0.04408 to 0.1134 | <0.0001 |
| Base vs. Post4 | 0.04955 | -0.01534 | 0.06489 | 0.00867 | 27 | 27 | 7.484 | 26 | 0.03867 to 0.09110 | <0.0001 |
| Base vs. Post5 | 0.04955 | 0.01999 | 0.02956 | 0.007947 | 27 | 27 | 3.719 | 26 | 0.005526 to 0.05359 | 0.0087 |
| Base vs. Post6 | 0.04955 | 0.03602 | 0.01353 | 0.008877 | 27 | 27 | 1.524 | 26 | -0.01331 to 0.04037 | >0.9999 |
| Base vs. Post7 | 0.04955 | 0.03715 | 0.01239 | 0.005516 | 27 | 27 | 2.247 | 26 | -0.004286 to 0.02907 | 0.3004 |
| Base vs. Post8 | 0.04955 | 0.04975 | -0.0002044 | 0.00441 | 27 | 27 | 0.0463 | 26 | -0.01354 to 0.01313 | >0.9999 |

**eTable 10: Bonferroni's multiple comparison test results comparing Pert1$_R$ to Pert10$_R$ for MoS$_{AP}$ in the older adults**

| Test details | Mean 1 | Mean 2 | Mean Diff. | SE of diff. | N1 | N2 | t | DF | 95.00% CI of diff. | Adjusted P Value |
|---|---|---|---|---|---|---|---|---|---|---|
| Pert1R - Pert10R | | | | | | | | | | |
| Base | 0.0496 | 0.04955 | 5.044E-05 | 0.001873 | 27 | 27 | 0.0269 | 26 | -0.005693 to 0.005794 | >0.9999 |
| Pre | 0.04576 | 0.04616 | -0.0003983 | 0.003603 | 27 | 27 | 0.1106 | 26 | -0.01145 to 0.01065 | >0.9999 |
| Post1 | 0.1117 | 0.01955 | 0.09217 | 0.01073 | 27 | 27 | 8.586 | 26 | 0.05925 to 0.1251 | <0.0001 |
| Post2 | 0.02267 | -0.03525 | 0.05792 | 0.01635 | 27 | 27 | 3.542 | 26 | 0.007766 to 0.1081 | 0.0152 |
| Post3 | -0.0008 | -0.0292 | 0.02842 | 0.01276 | 27 | 27 | 2.227 | 26 | -0.01072 to 0.06756 | 0.3485 |
| Post4 | -0.0279 | -0.01534 | -0.01252 | 0.012 | 27 | 27 | 1.043 | 26 | -0.04934 to 0.02429 | >0.9999 |
| Post5 | -0.0097 | 0.01999 | -0.02974 | 0.01126 | 27 | 27 | 2.642 | 26 | -0.06426 to 0.004782 | 0.1377 |
| Post6 | 0.0042 | 0.03602 | -0.03181 | 0.01121 | 27 | 27 | 2.838 | 26 | -0.06619 to 0.002567 | 0.0869 |
| Post7 | 0.01565 | 0.03715 | -0.0215 | 0.008091 | 27 | 27 | 2.658 | 26 | -0.04632 to 0.003310 | 0.1327 |
| Post8 | 0.02613 | 0.04975 | -0.02362 | 0.006687 | 27 | 27 | 3.533 | 26 | -0.04413 to -0.003115 | 0.0156 |

**eTable 11: Bonferroni's multiple comparison test results comparing recovery steps to baseline in Pert1$_R$ and Pert10$_R$ for BoS in the older adults**

| Test details | Mean 1 | Mean 2 | Mean Diff. | SE of diff. | N1 | N2 | t | DF | 95.00% CI of diff. | Adjusted P Value |
|---|---|---|---|---|---|---|---|---|---|---|
| Pert1$_R$ | | | | | | | | | | |
| Base vs. Pre | 0.6438 | 0.64 | 0.003826 | 0.004034 | 27 | 27 | 0.9484 | 26 | -0.008373 to 0.01603 | >0.9999 |
| Base vs. Post1 | 0.6438 | 0.7432 | -0.09937 | 0.01022 | 27 | 27 | 9.723 | 26 | -0.1303 to -0.06847 | <0.0001 |
| Base vs. Post2 | 0.6438 | 0.2436 | 0.4002 | 0.03247 | 27 | 27 | 12.33 | 26 | 0.3020 to 0.4984 | <0.0001 |
| Base vs. Post3 | 0.6438 | 0.5265 | 0.1173 | 0.02718 | 27 | 27 | 4.318 | 26 | 0.03516 to 0.1995 | 0.0018 |
| Base vs. Post4 | 0.6438 | 0.6003 | 0.04344 | 0.02796 | 27 | 27 | 1.554 | 26 | -0.04109 to 0.1280 | >0.9999 |
| Base vs. Post5 | 0.6438 | 0.6205 | 0.0233 | 0.01868 | 27 | 27 | 1.248 | 26 | -0.03317 to 0.07977 | >0.9999 |
| Base vs. Post6 | 0.6438 | 0.6482 | -0.004443 | 0.01213 | 27 | 27 | 0.3664 | 26 | -0.04111 to 0.03222 | >0.9999 |
| Base vs. Post7 | 0.6438 | 0.6157 | 0.02805 | 0.01354 | 27 | 27 | 2.071 | 26 | -0.01290 to 0.06899 | 0.4355 |
| Base vs. Post8 | 0.6438 | 0.629 | 0.01479 | 0.01053 | 27 | 27 | 1.404 | 26 | -0.01705 to 0.04663 | >0.9999 |
| Pert10$_R$ | | | | | | | | | | |
| Base vs. Pre | 0.6519 | 0.6503 | 0.001648 | 0.005721 | 27 | 27 | 0.288 | 26 | -0.01565 to 0.01895 | >0.9999 |
| Base vs. Post1 | 0.6519 | 0.7293 | -0.07739 | 0.01508 | 27 | 27 | 5.133 | 26 | -0.1230 to -0.03180 | 0.0002 |
| Base vs. Post2 | 0.6519 | 0.3814 | 0.2705 | 0.03636 | 27 | 27 | 7.441 | 26 | 0.1606 to 0.3805 | <0.0001 |
| Base vs. Post3 | 0.6519 | 0.5905 | 0.06141 | 0.01585 | 27 | 27 | 3.873 | 26 | 0.01347 to 0.1093 | 0.0059 |
| Base vs. Post4 | 0.6519 | 0.607 | 0.04488 | 0.02076 | 27 | 27 | 2.161 | 26 | -0.01791 to 0.1077 | 0.3605 |
| Base vs. Post5 | 0.6519 | 0.636 | 0.01587 | 0.01416 | 27 | 27 | 1.121 | 26 | -0.02695 to 0.05868 | >0.9999 |
| Base vs. Post6 | 0.6519 | 0.646 | 0.005856 | 0.009091 | 27 | 27 | 0.6442 | 26 | -0.02163 to 0.03334 | >0.9999 |
| Base vs. Post7 | 0.6519 | 0.6313 | 0.02063 | 0.01164 | 27 | 27 | 1.773 | 26 | -0.01455 to 0.05582 | 0.7912 |
| Base vs. Post8 | 0.6519 | 0.6456 | 0.006321 | 0.006719 | 27 | 27 | 0.9407 | 26 | -0.01400 to 0.02664 | >0.9999 |

**eTable 12: Bonferroni's multiple comparison test results comparing Pert1$_R$ to Pert10$_R$ for BoS in the older adults**

| Test details | Mean 1 | Mean 2 | Mean Diff. | SE of diff. | N1 | N2 | t | DF | 95.00% CI of diff. | Adjusted P Value |
|---|---|---|---|---|---|---|---|---|---|---|
| Pert1R - Pert10R | | | | | | | | | | |
| Base | 0.6438 | 0.6519 | -0.008115 | 0.004207 | 27 | 27 | 1.929 | 26 | -0.02102 to 0.004789 | 0.6476 |
| Pre | 0.64 | 0.6503 | -0.01029 | 0.006064 | 27 | 27 | 1.697 | 26 | -0.02889 to 0.008306 | >0.9999 |
| Post1 | 0.7432 | 0.7293 | 0.01386 | 0.0113 | 27 | 27 | 1.227 | 26 | -0.02078 to 0.04850 | >0.9999 |
| Post2 | 0.2436 | 0.3814 | -0.1378 | 0.04262 | 27 | 27 | 3.233 | 26 | -0.2685 to -0.007088 | 0.0332 |
| Post3 | 0.5265 | 0.5905 | -0.06405 | 0.02388 | 27 | 27 | 2.682 | 26 | -0.1373 to 0.009181 | 0.1253 |
| Post4 | 0.6003 | 0.607 | -0.00668 | 0.02648 | 27 | 27 | 0.2523 | 26 | -0.08788 to 0.07452 | >0.9999 |
| Post5 | 0.6205 | 0.636 | -0.01555 | 0.01339 | 27 | 27 | 1.161 | 26 | -0.05663 to 0.02553 | >0.9999 |
| Post6 | 0.6482 | 0.646 | 0.002185 | 0.01249 | 27 | 27 | 0.175 | 26 | -0.03611 to 0.04048 | >0.9999 |
| Post7 | 0.6157 | 0.6313 | -0.01553 | 0.01433 | 27 | 27 | 1.084 | 26 | -0.05948 to 0.02842 | >0.9999 |
| Post8 | 0.629 | 0.6456 | -0.01658 | 0.01148 | 27 | 27 | 1.444 | 26 | -0.05180 to 0.01863 | >0.9999 |

**eTable 13: Bonferroni's multiple comparison test results comparing recovery steps to baseline in Pert1$_R$ and Pert10$_R$ for $X_{CoM}$ in the older adults**

| Test details | Mean 1 | Mean 2 | Mean Diff. | SE of diff. | N1 | N2 | t | DF | 95.00% CI of diff. | Adjusted P Value |
|---|---|---|---|---|---|---|---|---|---|---|
| Pert1$_R$ | | | | | | | | | | |
| Base vs. Pre | 0.5942 | 0.5942 | -9.535E-06 | 0.005839 | 27 | 27 | 0.0016 | 26 | -0.01767 to 0.01765 | >0.9999 |
| Base vs. Post1 | 0.5942 | 0.6314 | -0.03724 | 0.01465 | 27 | 27 | 2.542 | 26 | -0.08154 to 0.007056 | 0.1559 |
| Base vs. Post2 | 0.5942 | 0.2209 | 0.3733 | 0.03143 | 27 | 27 | 11.88 | 26 | 0.2783 to 0.4683 | <0.0001 |
| Base vs. Post3 | 0.5942 | 0.5272 | 0.06696 | 0.02828 | 27 | 27 | 2.368 | 26 | -0.01856 to 0.1525 | 0.2307 |
| Base vs. Post4 | 0.5942 | 0.6282 | -0.03402 | 0.0273 | 27 | 27 | 1.246 | 26 | -0.1166 to 0.04853 | >0.9999 |
| Base vs. Post5 | 0.5942 | 0.6302 | -0.03604 | 0.02071 | 27 | 27 | 1.741 | 26 | -0.09866 to 0.02657 | 0.8421 |
| Base vs. Post6 | 0.5942 | 0.644 | -0.04984 | 0.0135 | 27 | 27 | 3.692 | 26 | -0.09065 to -0.009022 | 0.0093 |
| Base vs. Post7 | 0.5942 | 0.6001 | -0.0059 | 0.01252 | 27 | 27 | 0.4712 | 26 | -0.04376 to 0.03196 | >0.9999 |
| Base vs. Post8 | 0.5942 | 0.6029 | -0.00868 | 0.01323 | 27 | 27 | 0.656 | 26 | -0.04869 to 0.03133 | >0.9999 |
| Pert10$_R$ | | | | | | | | | | |
| Base vs. Pre | 0.6024 | 0.6041 | -0.001739 | 0.007199 | 27 | 27 | 0.2416 | 26 | -0.02351 to 0.02003 | >0.9999 |
| Base vs. Post1 | 0.6024 | 0.7097 | -0.1074 | 0.01619 | 27 | 27 | 6.635 | 26 | -0.1563 to -0.05844 | <0.0001 |
| Base vs. Post2 | 0.6024 | 0.4166 | 0.1857 | 0.0339 | 27 | 27 | 5.479 | 26 | 0.08324 to 0.2882 | <0.0001 |
| Base vs. Post3 | 0.6024 | 0.6197 | -0.01734 | 0.02256 | 27 | 27 | 0.7686 | 26 | -0.08557 to 0.05088 | >0.9999 |
| Base vs. Post4 | 0.6024 | 0.6224 | -0.02001 | 0.01829 | 27 | 27 | 1.094 | 26 | -0.07532 to 0.03530 | >0.9999 |
| Base vs. Post5 | 0.6024 | 0.616 | -0.01369 | 0.01932 | 27 | 27 | 0.7087 | 26 | -0.07210 to 0.04472 | >0.9999 |
| Base vs. Post6 | 0.6024 | 0.61 | -0.007673 | 0.01552 | 27 | 27 | 0.4944 | 26 | -0.05460 to 0.03925 | >0.9999 |
| Base vs. Post7 | 0.6024 | 0.5941 | 0.00824 | 0.01461 | 27 | 27 | 0.5641 | 26 | -0.03593 to 0.05241 | >0.9999 |
| Base vs. Post8 | 0.6024 | 0.5958 | 0.006525 | 0.008906 | 27 | 27 | 0.7327 | 26 | -0.02040 to 0.03345 | >0.9999 |

**eTable 14: Bonferroni's multiple comparison test results comparing Pert1$_R$ to Pert10$_R$ for $X_{CoM}$ in the older adults**

| Test details | Mean 1 | Mean 2 | Mean Diff. | SE of diff. | N1 | N2 | t | DF | 95.00% CI of diff. | Adjusted P Value |
|---|---|---|---|---|---|---|---|---|---|---|
| Pert1R - Pert10R | | | | | | | | | | |
| Base | 0.5942 | 0.6024 | -0.008165 | 0.004169 | 27 | 27 | 1.959 | 26 | -0.02095 to 0.004620 | 0.6095 |
| Pre | 0.5942 | 0.6041 | -0.009895 | 0.007249 | 27 | 27 | 1.365 | 26 | -0.03213 to 0.01234 | >0.9999 |
| Post1 | 0.6314 | 0.7097 | -0.07831 | 0.01123 | 27 | 27 | 6.976 | 26 | -0.1127 to -0.04388 | <0.0001 |
| Post2 | 0.2209 | 0.4166 | -0.1957 | 0.03769 | 27 | 27 | 5.193 | 26 | -0.3113 to -0.08013 | 0.0002 |
| Post3 | 0.5272 | 0.6197 | -0.09246 | 0.02854 | 27 | 27 | 3.24 | 26 | -0.1800 to -0.004932 | 0.0326 |
| Post4 | 0.6282 | 0.6224 | 0.005845 | 0.02627 | 27 | 27 | 0.2225 | 26 | -0.07471 to 0.08640 | >0.9999 |
| Post5 | 0.6302 | 0.616 | 0.01419 | 0.01757 | 27 | 27 | 0.8074 | 26 | -0.03970 to 0.06808 | >0.9999 |
| Post6 | 0.644 | 0.61 | 0.034 | 0.01895 | 27 | 27 | 1.794 | 26 | -0.02414 to 0.09213 | 0.8452 |
| Post7 | 0.6001 | 0.5941 | 0.005975 | 0.01438 | 27 | 27 | 0.4156 | 26 | -0.03812 to 0.05007 | >0.9999 |
| Post8 | 0.6029 | 0.5958 | 0.00704 | 0.01329 | 27 | 27 | 0.5298 | 26 | -0.03371 to 0.04779 | >0.9999 |

**eTable 15: Bonferroni's multiple comparison test results comparing recovery steps to baseline in Pert1$_R$ and Pert10$_R$ for MoS$_{ML}$ in the older adults**

| Test details | Mean 1 | Mean 2 | Mean Diff. | SE of diff. | N1 | N2 | t | DF | 95.00% CI of diff. | Adjusted P Value |
|---|---|---|---|---|---|---|---|---|---|---|
| Pert1$_R$ | | | | | | | | | | |
| Base vs. Pre | 0.01644 | 0.02095 | -0.004507 | 0.003311 | 27 | 27 | 1.361 | 26 | -0.01452 to 0.005503 | >0.9999 |
| Base vs. Post1 | 0.01644 | 0.005856 | 0.01058 | 0.008328 | 27 | 27 | 1.271 | 26 | -0.01460 to 0.03576 | >0.9999 |
| Base vs. Post2 | 0.01644 | 0.0454 | -0.02897 | 0.00871 | 27 | 27 | 3.326 | 26 | -0.05530 to -0.002629 | 0.0237 |
| Base vs. Post3 | 0.01644 | 0.03473 | -0.01829 | 0.009911 | 27 | 27 | 1.845 | 26 | -0.04826 to 0.01168 | 0.6876 |
| Base vs. Post4 | 0.01644 | 0.04115 | -0.02471 | 0.005466 | 27 | 27 | 4.521 | 26 | -0.04124 to -0.008186 | 0.0011 |
| Base vs. Post5 | 0.01644 | 0.02488 | -0.00844 | 0.00469 | 27 | 27 | 1.8 | 26 | -0.02262 to 0.005741 | 0.7519 |
| Base vs. Post6 | 0.01644 | 0.03278 | -0.01634 | 0.004258 | 27 | 27 | 3.837 | 26 | -0.02922 to -0.003465 | 0.0064 |
| Base vs. Post7 | 0.01644 | 0.02323 | -0.006792 | 0.004249 | 27 | 27 | 1.598 | 26 | -0.01964 to 0.006056 | >0.9999 |
| Base vs. Post8 | 0.01644 | 0.02829 | -0.01185 | 0.004343 | 27 | 27 | 2.73 | 26 | -0.02499 to 0.001278 | 0.101 |
| Pert10$_R$ | | | | | | | | | | |
| Base vs. Pre | 0.01969 | 0.02029 | -0.0005999 | 0.004008 | 27 | 27 | 0.1497 | 26 | -0.01272 to 0.01152 | >0.9999 |
| Base vs. Post1 | 0.01969 | 0.01298 | 0.006709 | 0.007452 | 27 | 27 | 0.9003 | 26 | -0.01582 to 0.02924 | >0.9999 |
| Base vs. Post2 | 0.01969 | 0.05607 | -0.03638 | 0.005344 | 27 | 27 | 6.808 | 26 | -0.05254 to -0.02022 | <0.0001 |
| Base vs. Post3 | 0.01969 | 0.02668 | -0.006992 | 0.006573 | 27 | 27 | 1.064 | 26 | -0.02687 to 0.01288 | >0.9999 |
| Base vs. Post4 | 0.01969 | 0.03991 | -0.02022 | 0.006154 | 27 | 27 | 3.286 | 26 | -0.03883 to -0.001612 | 0.0262 |
| Base vs. Post5 | 0.01969 | 0.0231 | -0.003408 | 0.003743 | 27 | 27 | 0.9105 | 26 | -0.01472 to 0.007909 | >0.9999 |
| Base vs. Post6 | 0.01969 | 0.03015 | -0.01046 | 0.004406 | 27 | 27 | 2.375 | 26 | -0.02379 to 0.002859 | 0.227 |
| Base vs. Post7 | 0.01969 | 0.02292 | -0.003227 | 0.004232 | 27 | 27 | 0.7626 | 26 | -0.01602 to 0.009568 | >0.9999 |
| Base vs. Post8 | 0.01969 | 0.02589 | -0.006201 | 0.002658 | 27 | 27 | 2.333 | 26 | -0.01424 to 0.001835 | 0.2487 |

**eTable 16: Bonferroni's multiple comparison test results comparing Pert1$_R$ to Pert10$_R$ for MoS$_{ML}$ in the older adults**

| Test details | Mean 1 | Mean 2 | Mean Diff. | SE of diff. | N1 | N2 | t | DF | 95.00% CI of diff. | Adjusted P Value |
|---|---|---|---|---|---|---|---|---|---|---|
| Pert1R - Pert10R | | | | | | | | | | |
| Base | 0.01644 | 0.01969 | -0.003249 | 0.001499 | 27 | 27 | 2.167 | 26 | -0.007847 to 0.001349 | 0.3955 |
| Pre | 0.02095 | 0.02029 | 0.0006582 | 0.003111 | 27 | 27 | 0.2115 | 26 | -0.008884 to 0.01020 | >0.9999 |
| Post1 | 0.00586 | 0.01298 | -0.007123 | 0.008802 | 27 | 27 | 0.8092 | 26 | -0.03412 to 0.01987 | >0.9999 |
| Post2 | 0.0454 | 0.05607 | -0.01067 | 0.009443 | 27 | 27 | 1.13 | 26 | -0.03963 to 0.01830 | >0.9999 |
| Post3 | 0.03473 | 0.02668 | 0.008048 | 0.01128 | 27 | 27 | 0.7132 | 26 | -0.02656 to 0.04266 | >0.9999 |
| Post4 | 0.04115 | 0.03991 | 0.001246 | 0.006638 | 27 | 27 | 0.1877 | 26 | -0.01911 to 0.02160 | >0.9999 |
| Post5 | 0.02488 | 0.0231 | 0.001783 | 0.005093 | 27 | 27 | 0.3502 | 26 | -0.01384 to 0.01740 | >0.9999 |
| Post6 | 0.03278 | 0.03015 | 0.002627 | 0.004233 | 27 | 27 | 0.6206 | 26 | -0.01036 to 0.01561 | >0.9999 |
| Post7 | 0.02323 | 0.02292 | 0.0003155 | 0.003709 | 27 | 27 | 0.0851 | 26 | -0.01106 to 0.01169 | >0.9999 |
| Post8 | 0.02829 | 0.02589 | 0.002405 | 0.004033 | 27 | 27 | 0.5962 | 26 | -0.009965 to 0.01477 | >0.9999 |

### eDiscussion

Regarding our analyses of $MoS_{ML}$, the results did not reveal any substantial differences with age. Both age groups showed a significant increase in $MoS_{ML}$ in some of the first few steps post-perturbation, but there was little change in magnitude of these values with repetition. This suggests that anteroposterior gait perturbations of this nature require some mediolateral balance control but can be accommodated similarly by young and older healthy adults. It is not possible to say, based on the current work, if different magnitudes or types of anteroposterior perturbations would show a similar lack of effect with age. Sideways falls are prevalent among older adults in long-term and many falls initially directed anteriorly may lead to a lateral ground contact [1]. Previous findings indicate that older age and falls history are related to lateral instability following anteroposterior perturbations to stance [2,3]. However, as the current study deals with perturbations during gait, stance perturbations are obviously not directly comparable to those used in the current paradigm. A limitation that should be noted is that while our reduced kinematic model has been previously validated for the $MoS_{AP}$ and its components [4], it was not validated for the $MoS_{ML}$. Various levels of agreement have been reported in previous studies of CoM position estimates between simplified marker sets and the full-body marker sets [5-8] and these differences can stem from differences in the marker model (e.g. with and without trunk), the analysed locomotion task (e.g. level walking, cutting manoeuvres), the participant groups (e.g. healthy young adults, stroke patients) and the gait velocity and experimental setup (e.g. overground and treadmill walking). Therefore, we would caution against drawing firm conclusions regarding our $MoS_{ML}$ results.
